# Using diffusion MRI data acquired with ultra-high gradients to improve tractography in routine-quality data

**DOI:** 10.1101/2021.06.28.450265

**Authors:** C. Maffei, C. Lee, M. Planich, M. Ramprasad, N. Ravi, D. Trainor, Z. Urban, M. Kim, R.J. Jones, A. Henin, S.G. Hofmann, D.A. Pizzagalli, R.P. Auerbach, J.D.E. Gabrieli, S. Whitfield-Gabrieli, D.N. Greve, S.N. Haber, A. Yendiki

**Affiliations:** Athinoula A. Martinos Center for Biomedical Imaging, Massachusetts General Hospital and Harvard Medical School, Charlestown, MA, USA; Massachusetts General Hospital and Harvard Medical School, Boston, MA, USA; Boston University, Boston, MA, USA; McLean Hospital and Harvard Medical School, Belmont, MA, USA; Columbia University, New York, NY, USA; Massachusetts Institute of Technology, Cambridge, MA, USA; Northeastern University, Boston, MA, USA; Department of Pharmacology and Physiology, University of Rochester School of Medicine, Rochester, NY, USA

**Keywords:** Diffusion MRI, Tractography, White matter pathways, Neuroanatomy, Anatomical priors

## Abstract

The development of scanners with ultra-high gradients, spearheaded by the Human Connectome Project, has led to dramatic improvements in the spatial, angular, and diffusion resolution that is feasible for *in vivo* diffusion MRI acquisitions. The improved quality of the data can be exploited to achieve higher accuracy in the inference of both microstructural and macrostructural anatomy. However, such high-quality data can only be acquired on a handful of Connectom MRI scanners worldwide, while remaining prohibitive in clinical settings because of the constraints imposed by hardware and scanning time. In this study, we first update the classical protocols for tractography-based, manual annotation of major white-matter pathways, to adapt them to the much greater volume and variability of the streamlines that can be produced from today’s state-of-the-art diffusion MRI data. We then use these protocols to annotate 42 major pathways manually in data from a Connectom scanner. Finally, we show that, when we use these manually annotated pathways as training data for global probabilistic tractography with anatomical neighborhood priors, we can perform highly accurate, automated reconstruction of the same pathways in much lower-quality, more widely available diffusion MRI data. The outcomes of this work include both a new, comprehensive atlas of WM pathways from Connectom data, and an updated version of our tractography toolbox, TRActs Constrained by UnderLying Anatomy (TRACULA), which is trained on data from this atlas. Both the atlas and TRACULA are distributed publicly as part of FreeSurfer. We present the first comprehensive comparison of TRACULA to the more conventional, multi-region-of-interest approach to automated tractography, and the first demonstration of training TRACULA on high-quality, Connectom data to benefit studies that use more modest acquisition protocols.

## 1. Introduction

Diffusion MRI (dMRI) tractography allows us to investigate the connectional anatomy of the human brain *in vivo* and non-invasively. One of its applications is the delineation of white-matter (WM) bundles that are known from the anatomical literature, with the goal of studying their macro-and micro-structural properties in both healthy and clinical populations.

Different methods have been proposed for extracting these bundles from whole-brain tractograms. The majority of these methods follow the multi-region of interest (multi-ROI) approach. Multi-ROI methods can be manual or automated. In the former case, ROIs are hand-drawn in individual dMRI space by an operator (Catani and Thiebaut de Schotten 2008; Wakana et al. 2007; Thiebaut de Schotten et al. 2011). For each WM bundle of interest, a set of *a priori* rules define which ROIs the bundle does or does not go through. The rules are applied to tractography streamlines obtained from each individual’s dMRI data, and the ROIs are refined manually to obtain bundles that match the anatomical literature as closely as possible. This manual procedure is tailored to each individual subject, and therefore has the potential to achieve high anatomical accuracy, to the extent that the initial streamlines obtained from the subject’s dMRI data are accurate. However, it is time-intensive and requires extensive prior anatomical knowledge on the part of the operator, limiting reproducibility and applicability to large datasets (Rheault et al. 2020). Automated multi-ROI methods follow a similar approach, but derive the ROIs either from atlases (Clayden et al. 2009; Yeatman et al. 2012; Groot et al. 2013; W. Zhang et al. 2008) or from automated, subject-specific, anatomical segmentations (Wassermann et al. 2013). This is faster than the manual approach and not operator-dependent. However, methods that rely on accurate registration of each individual to an atlas may be sensitive to individual anatomical variability. Importantly, both manual and automated multi-ROI methods are applied to tractography streamlines as a post-processing step. As a result, their accuracy is intrinsically limited by the quality of those streamlines, and therefore by the quality of the individual dMRI data. An alternative family of bundle segmentation methods relies on clustering algorithms, which group whole-brain tractography streamlines into clusters based on their similarity (O’Donnell et al. 2007; Visser et al. 2011; Garyfallidis et al. 2012; Ros et al. 2013; Siless et al. 2018). Each cluster of streamlines can then be labeled as a specific WM bundle, either based on its similarity to manually labeled bundles, or based on multi-ROI rules (Wasserman et al. 2010; Guevara et al. 2012; Garyfallidis et al. 2018; F. Zhang et al. 2018).

All of the above methods perform *post hoc* classification of tractography streamlines. If a subject’s tractogram does not contain any streamlines from a certain WM bundle, these methods will not be able to recover this bundle. Previous studies have shown that the precision and reliability of tractography are largely influenced by image quality and hence by the acquisition protocol (Jbabdi and Johansen-Berg 2011; Vos et al. 2012; Calabrese et al. 2014; C. Maffei, Sarubbo, and Jovicich 2019). The technical advances spearheaded by the Human Connectome Project (HCP) led to MRI systems with ultra-high gradients, which can achieve high diffusion weighting (b-value) without loss of signal-to-noise ratio, as well as accelerated MRI sequences that enable high angular and spatial resolution (Setsompop et al. 2013). However, dMRI data acquired in clinical settings typically have much lower quality, due to MRI hardware limitations and scan time constraints. This limits the accuracy of tractography, especially in bundles that are challenging because of their anatomical location, size or shape. The multi-ROI methods described above cannot address this. Even if the ROIs are defined on an atlas obtained from high-quality data, they cannot improve the reconstruction of WM bundles in individual data collected with poorer signal-to-noise ratio, spatial or angular resolution.

In this study we demonstrate how WM bundles labeled manually in high-quality data can be used to ensure accurate, automated reconstruction of the same bundles in routine-quality data. First, we describe a protocol for the manual dissection of 42 WM bundles from high-quality, high-b data collected on a Connectom scanner by the HCP. These data allow us to generate a much more detailed and accurate definition of the major bundles of the human brain than what would be possible from routine-quality data. Our virtual dissection protocols are more detailed than previously proposed ones (Wakana et al. 2007), to handle the much greater volume and variability of the streamlines produced by today’s state-of-the-art data acquisition, orientation modeling, and tractography methods.

Second, we use these manually dissected WM bundles as a new training dataset for TRACULA (TRActs Constrained by UnderLying Anatomy) (Yendiki et al. 2011). In contrast to multi-ROI or clustering-based methods for bundle reconstruction, TRACULA does not operate on tractography streamlines as a post-processing step. Instead, it incorporates prior information on WM anatomy in the tractography step itself. This is done via a Bayesian framework for global tractography that incorporates prior probabilities on the anatomical neighborhood of WM bundles. Here we demonstrate that, when these prior probabilities are computed from high-quality training data, TRACULA can reconstruct the same bundles in routine-quality data with high anatomical accuracy. Specifically, we train TRACULA on bundles labeled manually from HCP data with a maximum b-value of 10,000 *s*/*mm*^2^, and use it to reconstruct the same 42 bundles from data acquired with a b-value of 1,000 *s*/*mm*^2^. We compare these reconstructions to those obtained by an automated multi-ROI approach. We show that TRACULA achieves overall higher accuracy and reliability.

The contribution of this work is twofold: *(i)* an updated set of protocols for manual dissection of 42 WM bundles that are appropriate for tractograms obtained from state-of-the-art Connectom data and *(ii)* a demonstration of automated tractography that can achieve a form of “quality transfer” from Connectom data to more routine, clinical-quality data. Both the manually labeled tracts, and the refactored version of TRACULA that uses them as training data, are included in FreeSurfer 7.2 (https://github.com/freesurfer/freesurfer/tree/fs-7.2-beta). Extensive documentation and tutorials are available on the FreeSurfer wiki (https://surfer.nmr.mgh.harvard.edu/fswiki/Tracula). Visualizations of the 42 manually annotated WM bundles, as well as along-tract profiles of microstructural measures on these bundles, are available at: https://dmri.mgh.harvard.edu/tract-atlas/.

## 2. Methods

### 2.1 Overview

We used state-of-the-art tractography techniques on the b_max_=10,000 *s*/*mm*^2^ HCP data to produce high-quality, whole-brain tractograms. We applied a manual, multi-ROI approach to delineate a set of 42 WM bundles from these tractograms. We then used the manually annotations to inform two methods (TRACULA and multi-ROI) for reconstructing the same bundles automatically from the b=1,000 *s*/*mm*^2^ data of the same subjects. We quantified the accuracy of each method by computing the distance of the bundles that were reconstructed automatically on the b=1,000 *s*/*mm*^2^ data from those that were annotated manually on the b_max_=10,000 *s*/*mm*^2^ data of the same subject. We also assessed the test-retest reliability of along-tract microstructural measures obtained from the automatically reconstructed bundles, either with TRACULA or with the multi-ROI method. Finally, we used this updated version of TRACULA to study associations between WM microstructure and psychopathology in a larger, independent dataset.

### 2.2 Data

The manual annotation used diffusion and structural MRI data of 16 healthy adult subjects provided by the MGH-USC HCP. The dMRI data include 512 diffusion-weighted (DW) volumes (b-values= 1,000/3,000/5,000/10,000 *s*/*mm*^2^) and 40 non-DW volumes (b=0) with 1.5 *mm* isotropic spatial resolution (Fan et al. 2015). The structural (T1-weighted) data were acquired with a multi-echo magnetization-prepared rapid acquisition gradient echo (MEMPRAGE) sequence at 1 *mm* isotropic resolution.

### 2.3 Image analysis

#### 2.3.1 Structural MRI

Cortical parcellations and subcortical segmentations were obtained for each subject using FreeSurfer (Dale, Fischl, and Sereno 1999; Fischl, Sereno, and Dale 1999, Fischl et al. 2002; Fischl et al. 2004). Segmentations of the thalamic nuclei and hypothalamic subunits were also obtained for each subject (Iglesias et al. 2015, 2018).

#### 2.3.2 Diffusion MRI

Diffusion data were denoised (Veraart et al. 2016) and corrected for gradient nonlinearity distortions (Glasser et al., 2013; Jovicich et al., 2006). Data were then corrected for head motion and eddy-current artifacts using *eddy* in FSL 6.0.3 (Andersson et al. 2016a, Andersson et al. 2016b). For each subject, we obtained whole-brain probabilistic tractograms using two methods: constrained spherical deconvolution (CSD) (Tax et al. 2014) on the *b* = 10,000 *s*/*mm*^2^ shell only (step-size: 0.5 *mm*, angle-threshold: 30^∘^, 10 seeds/voxel in white matter mask) in DIPY (Garyfallidis et al. 2014) and multi-shell multi-tissue CSD (MSMT-CSD) (Jeurissen et al. 2014) on all four shells (step-size: 0.75 *mm*, angle-threshold: 45^∘^, 50 seeds/voxel in white matter mask) in MRtrix3 (Tournier et al. 2012). We used partial volume masks of WM, gray matter (GM), and cerebrospinal fluid (CSF) to constrain the tractography results (e.g., ensure that streamlines terminate at the GM-WM interface) (Smith et al. 2012). We chose these two streamline tractography approaches empirically, after testing several state-of-the-art, publicly available methods, as they yielded sharp orientation distribution functions in fiber-crossing regions and in regions with partial voluming, respectively.

### 2.4 Manual labeling in high-quality data

We dissected 42 WM pathways manually in Trackvis (v.0.6.1; http://www.trackvis.org). For each tract, we defined a combination of inclusion and exclusion ROIs in the space of each individual subject. We derived protocols for the placement of these ROIs based on the anatomical literature, as detailed in the following sections. Streamlines from an individual’s whole-brain tractogram (described in the previous section) were retained if they passed through all inclusion ROIs and discarded if they passed through any of the exclusion ROIs defined for a specific bundle. Any FreeSurfer cortical ROIs that were used for the manual dissection came from the Desikan-Killiany parcellation (Desikan et al. 2006) and were grown 5 *mm* into the WM, along the normal vector of the cortical surface. The FreeSurfer corpus callosum (CC) ROIs, wherever used, came from the subcortical segmentation and covered only the section of the CC between the two hemispheres, along the midline. All projection and association pathways were dissected in the left and right hemisphere, denoted in the following as LH and RH, respectively. Each pathway was labeled by a single rater and then checked by CM for correctness and consistency with neighboring pathways.

#### 2.4.1 Commissural pathways

The manual labeling protocol for these pathways is illustrated in Fig. 1.

**Fig. 1.**
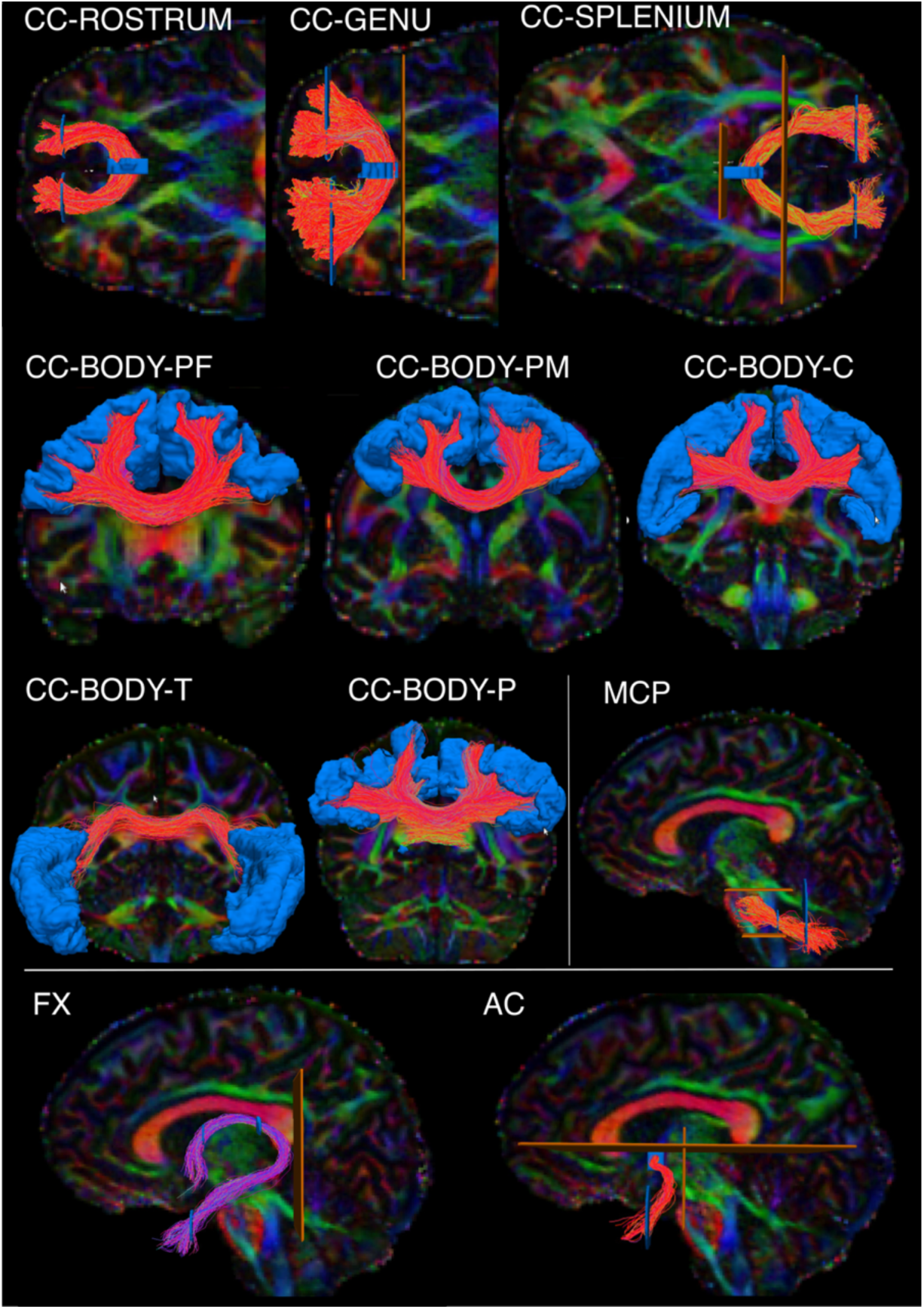
Manual labeling: commissural pathways. The figure shows the manual labeling protocols for the commissural pathways in one representative subject. Inclusion ROIs are shown in blue, exclusion ROIs in orange. Tracts are shown on color-coded FA maps. CC: Corpus callosum. It is subdivided into the rostrum, genu, splenium, and body. The body is further subdivided into prefrontal (BODY-PF), premotor (BODY-PM), central (BODY-C), temporal (BODY-T), and parietal (BODY-P) components. MCP: middle cerebellar peduncle. FX: fornix. AC: anterior commissure.

##### The Anterior Commissure (AC)

The AC was defined as a fiber bundle running transversely between the anterior part of the bilateral temporal lobes and situated below the fornix medially and the uncinate fascicle laterally (J. Schmahmann and Pandya 2006). We used color-coded fractional anisotropy (FA) maps to draw a first inclusion ROI around the left-right oriented region in front of the anterior columns of the fornix (sagittal view). Although it has been suggested that the AC also includes posterior projections to the occipital lobe (Turner, Mishkin, and Knapp 1979), we decided to include only the anterior limb of the AC terminating in the WM of the temporal poles, as this is what is most commonly referred to as the AC (Catani and Mesulam 2008; Lawes et al. 2008). Two more inclusion ROIs were thus drawn to encompass the WM of the temporal pole in each hemisphere. A coronal ROI was used to exclude the posterior projections, and two sagittal ROIs were used to exclude the most lateral fibers of the AC adjacent to the external capsule.

##### The Corpus Callosum (CC)

*Genu:* The FreeSurfer segmentation label of the mid-anterior CC was used to select the streamlines of the genu. A second and third ROI including medial and lateral regions of the frontal lobe were used to include only frontal projections in both hemispheres and discard spurious streamlines.

*Rostrum:* The FreeSurfer segmentation label of the anterior CC was used to select the streamlines of the rostrum. A second and third ROI were used to include only streamlines terminating in the orbital regions of the frontal cortex in each hemisphere.

*Splenium:* The splenium was defined as connecting parietal and occipital cortices. Streamlines projecting to the temporal lobe were not included. The FreeSurfer regions of the posterior and mid-posterior CC were used to select the streamlines of the splenium. A second and third ROI encompassing the occipital and parietal WM were used to include only the streamlines projecting posteriorly in each hemisphere.

*Body*: The inclusion ROIs of the genu, rostrum, and splenium in the frontal and occipital WM were used as exclusion ROIs, to isolate the body of the CC from all other streamlines crossing the FreeSurfer midline CC labels. Given the topographic organization of the CC, we further subdivided the body into 5 sections, based on the cortical terminations of the streamlines. The temporal section (BODY-T) included terminations in the FreeSurfer regions: superior temporal, middle temporal, inferior temporal, transverse temporal, and banks of the superior temporal sulcus. The parietal section (BODY-P) included terminations in regions: superior parietal, supramarginal, and precuneus. The central section (BODY-C) included terminations in regions: precentral, postcentral, and paracentral. Subdividing the remaining (prefrontal and premotor) terminations of the body required subdividing the superior frontal parcellation label, which is large and spans both of those termination areas. We used a boundary from a previously proposed, publicly available parcellation scheme, which translated anatomical definitions of cytoarchitectonic regions of the frontal cortex from Petrides et al. 2012 to the fsaverage cortical surface (Tang et al. 2019). We mapped that parcellation from the fsaverage surface to the individual surface of each training subject using the inverse of the FreeSurfer spherical morph. We used the boundary that separated areas 6, 8, and 44 from areas 9, 46, and 45 in that parcellation to subdivide the individual superior frontal label from FreeSurfer into a caudal and a rostral parcel. We then defined a premotor section of the body of the CC (BODY-PM) that included terminations in the caudal subdivision of the superior frontal label or in the FreeSurfer caudal middle frontal label. Finally, we defined a prefrontal section of the body of the CC (BODY-PF) that included terminations in the rostral subdivision of the superior frontal label or in the FreeSurfer rostral middle frontal label.

*The Fornix (FX).* The FX was defined as streamlines surrounding the thalamus, directly adjacent to the medial half of its superior and posterior surfaces (Pascalau et al. 2018) and connecting the hippocampal formation (specifically CA1, CA3, and fimbria) with the anterior thalamic nuclei, the mammillary bodies, the medial septal nucleus, and the basal forebrain (Poletti and Creswell 1977; Christiansen et al. 2017). A first inclusion ROI was placed on the coronal plane, inferior to the body of the CC, to outline the fornix body. A second inclusion ROI was then placed inferior and lateral to the hippocampus, where the fornix terminates. The subnuclei of the hippocampus (CA1, CA3, fimbria) (Iglesias et al. 2015) were used to confirm the correct terminations of the fornix. The tract was refined by placing two more inclusion ROIs anterior to the splenium of CC on a coronal slice to encompass each respective crus of the fornix. One exclusion ROI was then placed posterior to the crus to discard spurious streamlines.

#### 2.4.2 Projection pathways

The manual labeling protocol for these pathways is illustrated in Fig. 2.

**Fig. 2.**
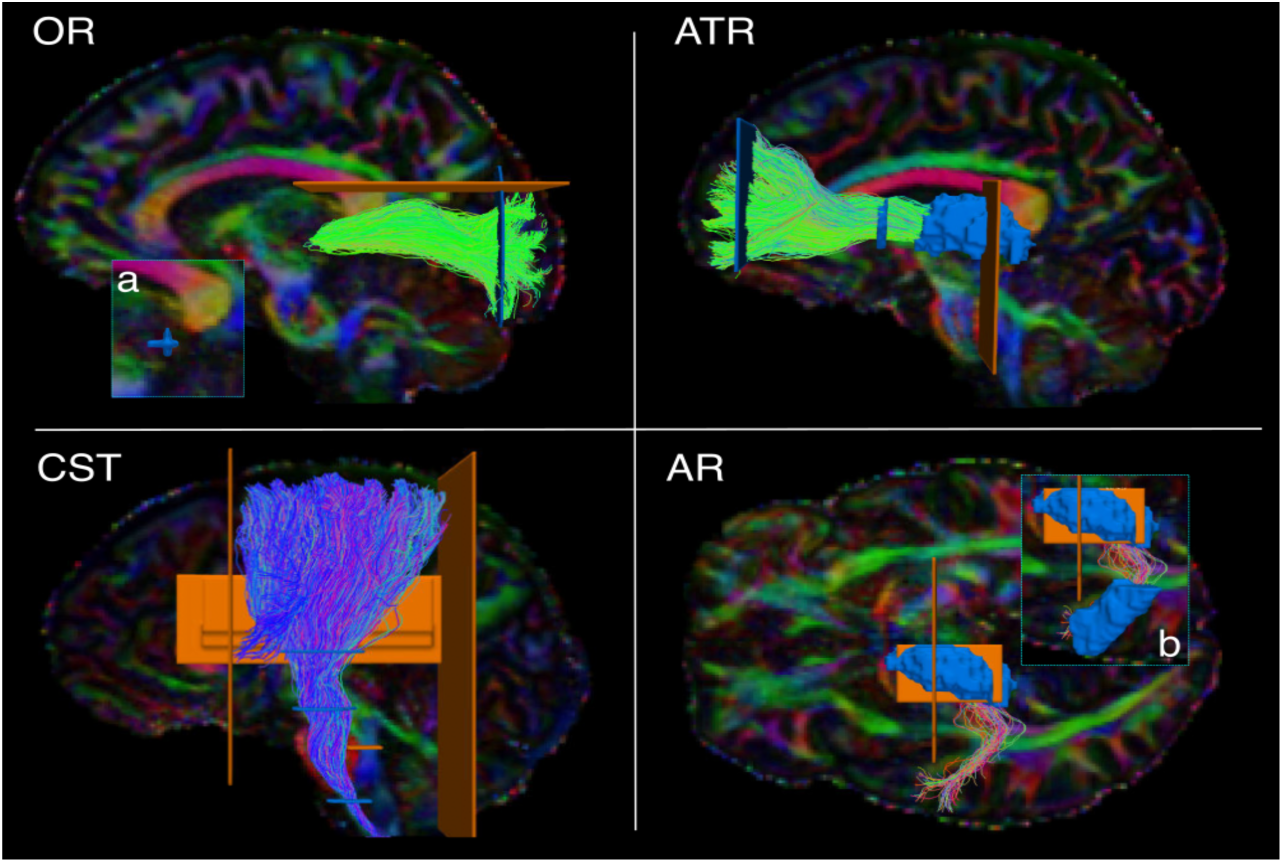
Manual labeling: projection pathways. The figure shows the manual labeling protocols for the projection pathways in one representative subject. Inclusion ROIs are shown in blue, exclusion ROIs in orange. Tracts are shown on color-coded FA maps. OR: optic radiation. ATR: anterior thalamic radiation. CST: cortico-spinal tract. AR: acoustic radiation. a) zoom-in showing ROI on the lateral geniculate nucleus of the thalamus. b) zoom-in showing ROI on Heschl’s gyrus.

##### The Acoustic Radiation (AR)

The AR was defined as fibers originating in posterior thalamus, where the medial geniculate nucleus (MGN) is located, and terminating on the transverse temporal gyrus of Heschl (HG) in the posterior portion of the superior temporal gyrus (STG) (Bürgel et al. 2006; Rademacher, Bürgel, and Zilles 2002; Chiara Maffei et al. 2018). The FreeSurfer segmentation label of the entire thalamus was used as a first inclusion ROI, and a second inclusion ROI was manually drawn to encompass the GM and WM of the HG as previously described (C. Maffei, Sarubbo, and Jovicich 2019).

##### The Anterior Thalamic Radiation (ATR)

The ATR was defined as fibers originating in the anterior and medial thalamus, passing through the anterior limb of the internal capsule (ALIC), and connecting to the prefrontal cortex (Wakana et al. 2007). The Freesurfer segmentation label of the entire thalamus was used as the first inclusion ROI. A second inclusion ROI was drawn on a coronal slice to encompass the prefrontal WM of the superior and middle frontal gyrus. A third inclusion ROI was drawn on the ALIC on a coronal slice. An exclusion ROI was placed on the midline (sagittal plane) to remove streamlines crossing to the contralateral hemisphere through the CC.

##### The Cortico-Spinal Tract (CST)

The CST was defined as streamlines passing through the midbrain, the medulla oblongata, and the internal capsule (first, second, third inclusion ROI, respectively). We retained its terminations in the precentral and postcentral gyri, as well as the posterior third of the superior frontal gyrus, corresponding to the supplementary motor area (SMA) (Chenot et al. 2019). Two coronal exclusion ROIs were placed to discard streamlines projecting too anteriorly or posteriorly: one posterior to the postcentral sulcus, and one anterior to the SMA. Additional exclusion ROIs were drawn on the midline (sagittal plane) and the tegmental tract (axial plane).

##### The Optic Radiation (OR)

The OR was defined as connecting the thalamus and the occipital cortex (Kammen et al. 2016; Sarubbo et al. 2015). The whole thalamus as segmented in FreeSurfer was used as a first inclusion ROI. A second inclusion ROI (coronal plane) was used to encompass the WM of the occipital lobe. An exclusion ROI (coronal plane) was used to discard the posterior projections of the CC. Another exclusion ROI was drawn on the axial plane to discard streamlines projecting too superiorly.

#### 2.4.3 Association pathways

The manual labeling protocol for these pathways is illustrated in Fig. 3.

**Fig. 3.**
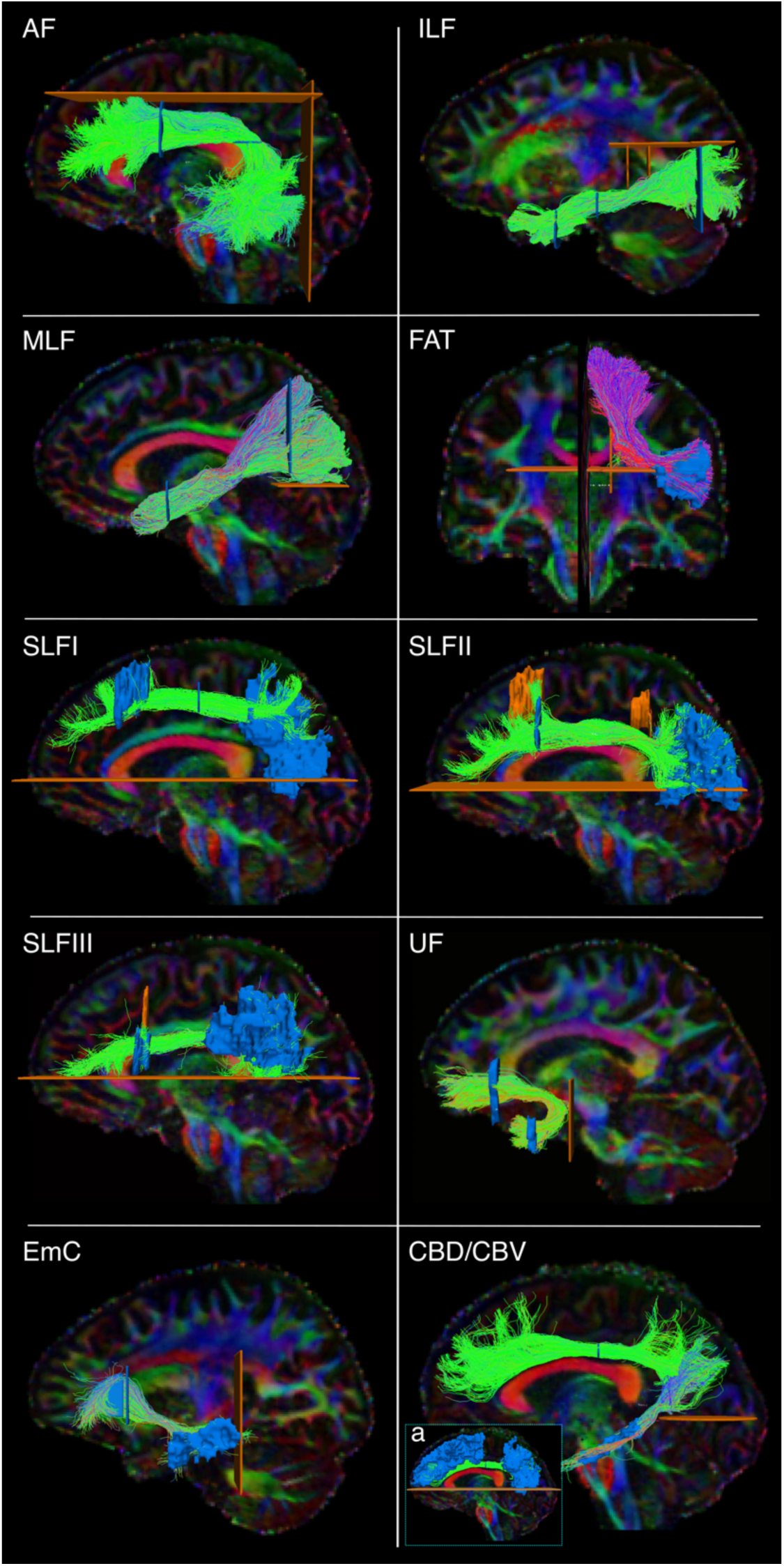
Manual labeling: association pathways. The figure shows the manual labeling protocols for the association pathways in one representative subject. Inclusion ROIs are shown in blue, exclusion ROIs in orange. Tracts are shown on color-coded FA maps. AF: arcuate fasciculus. ILF: inferior longitudinal fasciculus. MLF: middle longitudinal fasciculus. FAT: frontal aslant tract. SLF: superior longitudinal fasciculus. UF: uncinate fasciculus. EmC: extreme capsule. CBD/CBV: dorsal and ventral part of the cingulum bundle. a) inclusion and exclusion ROIs for the CBD.

##### The Arcuate Fasciculus (AF)

The AF was defined as the long, direct connections arching around the Sylvian fissure and connecting temporal (inferior, middle, and superior temporal gyri) and frontal regions (Catani, Jones, and Ffytche 2005; Lawes et al. 2008; J. D. Schmahmann et al. 2007; Makris et al. 2005; Fernández-Miranda et al. 2015). A first inclusion ROI was drawn on 3 consecutive axial slices at the level of the main body of the CC (medial boundary: line between arcuate and corona radiata; lateral boundary: postcentral sulcus; anterior boundary: precentral sulcus; posterior boundary: intraparietal sulcus). A second inclusion ROI was placed on a coronal slice at the level of the precentral sulcus (medial boundary: lateral ventricle; lateral/ventral/dorsal boundary: GM around Sylvian fissure and parietal lobe sulci) (Catani and Mesulam 2008). One exclusion ROI was drawn on a sagittal slice just lateral to the corona radiata, to remove erroneously crossing streamlines to the contralateral hemisphere. Two additional exclusion ROIs were placed superior and posterior to the AF to remove spurious streamlines.

##### The Cingulum Bundle (CB)

The CB was defined as a long associative bundle running in the WM adjacent to the cingulate gyrus (CG), arching around the splenium of the CC at the level of the cingulate isthmus, and terminating at the parahippocampal gyrus (J. Schmahmann and Pandya 2006; Lawes et al. 2008). To isolate the CB streamlines, a first ROI was drawn to include the anterior-posterior oriented regions superior to the CC as identified on coronal color-coded FA maps. We then subdivided the CB in two sub-bundles (Wakana et al. 2007; Jones, Knösche, and Turner 2013): a dorsal component running in the CG (CBD) and a ventral component running in the parahippocampal gyrus (CBV). We defined the CBD as connecting the anterior CG and the superior frontal gyrus (SFG) with parietal WM superior to the splenium of the CC, and the CBV as connecting these superior regions with the parahipopcampal gyrus. One exclusion ROI was placed on one axial slice inferior to the splenium of the CC to exclude ventral streamlines from the CBD (Fig. 3a), and one on one axial slice inferior and posterior to the splenium of the CC for the CBV.

##### The Extreme Capsule (EmC)

The EmC was defined as streamlines connecting the frontal and temporal regions, and located lateral to the uncinate fasciculus (UF) (Heide et al 2013). A first hand-drawn inclusion ROI was placed in the SFG to encompass most of the WM Brodmann’s areas 9 and 10 (Mars et al. 2016; Makris et al. 2009). This ROI was placed on the sagittal plane to make sure to distinguish EmC streamlines projecting laterally from UF streamlines projecting anteriorly (see below for UF dissection protocol). A second hand-drawn inclusion ROI was placed in the MTG. An exclusion ROI was located on the coronal plane posterior to the STG. A large exclusion ROI was placed along the midline of the brain.

##### The Frontal Aslant Tract (FAT)

The FAT was defined as streamlines connecting the posterior inferior frontal gyrus (IFG), pars opercularis, and medial aspects of the SFG, namely the pre-SMA and SMA (J. Schmahmann and Pandya 2006; Dick et al. 2019; Lawes et al. 2008). Exclusion ROIs were placed on a coronal slice posterior to the SMA and anterior to the pre-SMA, on the sagittal plane to exclude streamlines entering the CC, and on the axial plane to exclude artefactual streamlines projecting inferior.

##### The Inferior Longitudinal Fasciculus (ILF)

The ILF was defined as streamlines connecting superior, middle, inferior occipital gyri, and the fusiform and lingual gyri to the inferior and middle temporal gyri and the temporal pole (Latini et al. 2017). A first inclusion ROI was placed on a coronal slice, at the level of the precentral sulcus, to outline the temporal lobe, excluding the superior temporal sulcus. A second inclusion ROI was placed posterior to the CBD on a coronal slice to encompass the occipital WM. One exclusion ROI was placed superiorly (axial plane) to discard parietal connections, and one medially to the ILF (sagittal plane) to discard spurious streamlines.

##### The Middle Longitudinal Fasciculus (MLF)

The MLF was defined as streamlines connecting the superior and middle anterior temporal gyri and the temporal pole with the superior and inferior parietal cortex, coursing medial to the AF and superior to the ILF (Menjot De Champfleur et al. 2013; N. Makris et al. 2013; J. Schmahmann and Pandya 2006; Maldonado et al. 2013). A first inclusion ROI was placed on a coronal slice at the level of the precentral sulcus, to outline the superior temporal lobe. A second inclusion ROI was placed posterior to the CBD on a coronal slice to include both the superior and inferior parietal WM. An exclusion ROI was placed on the axial plane at the level of the parieto-occipital sulcus to discard streamlines going into the occipital lobe.

##### The Superior Longitudinal Fasciculus (SLF)

We dissected three SLF branches following definitions from the anatomical literature (J. D. Schmahmann et al. 2007; Hecht et al. 2015; Howells et al. 2018). SLF1: We placed one inclusion ROI in the superior frontal gyrus and one encompassing the WM posterior to the posterior central gyrus and dorsal to the cingulate sulcus. SLF2: We placed one inclusion ROI in the caudal part of the middle frontal gyrus and one in the WM of the inferior parietal lobe (Thiebaut de Schotten et al. 2011; Makris et al. 2009). SLF3: We placed one inclusion ROI in the posterior inferior frontal gyrus and one in the anterior supramarginal gyrus (J. D. Schmahmann et al. 2007; Hecht et al. 2015; Howells et al. 2018). For all three bundles, we used mid-sagittal and temporal exclusion ROIs.

##### The Uncinate Fasciculus (UF)

The UF was defined as streamlines connecting the anterior temporal pole and anterior middle temporal gyrus (MTG) with the medial and orbital prefrontal cortex (J. Schmahmann and Pandya 2006; Catani and Mesulam 2008). These streamlines were identified as medial and inferior to the EmC. The first inclusion ROI was drawn on four consecutive coronal slices in the temporal lobe, to encompass the WM of the MTG and temporal pole. A second inclusion ROI was drawn in the frontal lobe on four consecutive coronal slices on the WM of the medial orbito-frontal cortex. The subgenual WM was considered the upper limit of this ROI. Exclusion ROIs were placed on the mid-sagittal slice between the two hemispheres and directly posterior to the stem of the UF to exclude erroneous streamlines. We ensured that the relative position of the UF with respect to the EmC was accurate in each subject by labeling these two tracts jointly.

### 2.5 Automated reconstruction in routine-quality data

The bundles that were labeled manually in the b_max_ =10,000 *s*/*mm*^2^ data, were also reconstructed automatically in the b=1000 *s*/*mm*^2^ data of the same subjects (Fig. 4). The b=1000 *s*/*mm*^2^ shell comprised 64 out of the 512 DW volumes. We compared two approaches to automated reconstruction: (i) TRACULA, where we used the manually labeled bundles from the b_max_ = 10,000 *s*/*mm*^2^ data to compute prior probabilities on the anatomical neighborhood of each bundle and incorporated them in a Bayesian framework global probabilistic tractography, and (ii) Multi-ROI, where we used the group-averaged ROIs and inclusion/exclusion rules from the manual labeling as post-hoc constraints for local probabilistic tractography. We evaluated both approaches in a leave-one-out scheme, where the automated reconstruction in each subject used the manually labeled bundles or the labeling ROIs from the other 15 subjects.

**Fig. 4.**
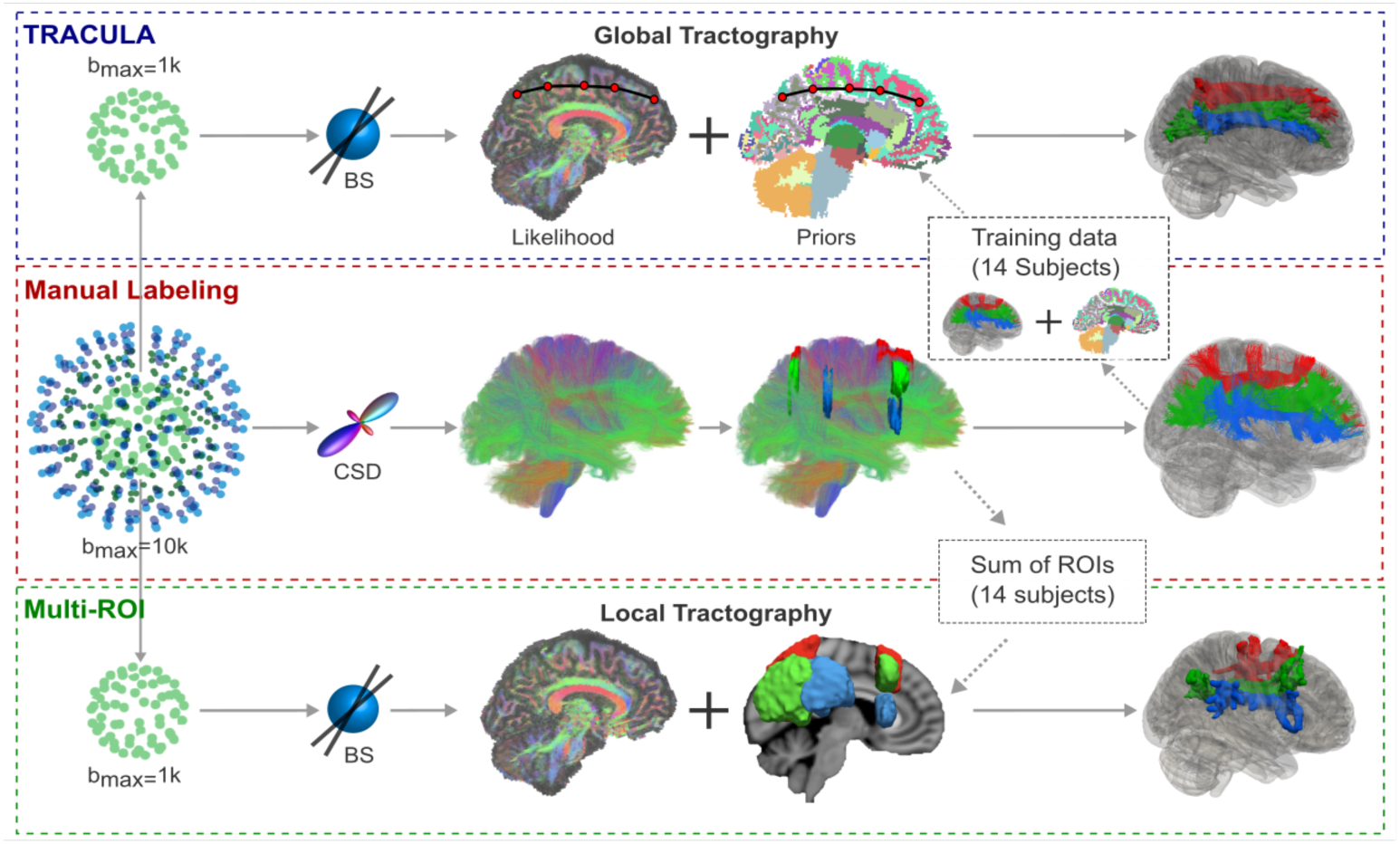
Overview of tractography methods. From the four-shell MGH-USC HCP data, the b=10,000 *s*/*mm*^2^ and b=1000 *s*/*mm*^2^ shells were extracted. Orientations were reconstructed with constrained spherical deconvolution (CSD) from the b=10,000*s*/*mm*^2^ shell and with multi-shell multi-tissue CSD (MSMT-CSD) from all four shells. Streamline tractography was performed with these two approaches and used to annotate 42 tracts manually in 16 subjects. The lower shell (b=1000 *s*/*mm*^2^, 64 directions) was used to reconstruct the same tracts automatically, with TRACULA or with a multi-ROI approach. For TRACULA, anatomical priors for each subject were obtained from the other 15 subjects and global probabilistic tractography was performed. For the multi-ROI approach, inclusion and exclusion masks were obtained from summing the manually defined ROIs of the other 15 subjects in template space. Local probabilistic tractography was constrained by these ROIs. The same ball-and-stick (BS) diffusion model was used for both TRACULA and the multi-ROI approach.

#### 2.5.1 TRACULA

##### Training data

The manual labeling procedure of section 2.4 produced a total of 2.29 million streamlines over all 42 bundles and 16 training subjects, covering 82% of all cerebral and cerebellar WM voxels. (In comparison, the manually labeled training set used in previous versions of our software included a total of 0.15 million streamlines from 18 bundles, which had been labeled in much lower-quality data and covered 18% of WM voxels.) This required us to refactor the TRACULA code base extensively to be able to handle a much larger training set than before. In this new, refactored version, many of the operations involved in computing the anatomical neighborhood priors, which were previously computed on the fly, are now precomputed and stored with the publicly distributed training data.

In addition, the densest of the manually labeled bundles, *e.g.,* most subdivisions of the CC, included a large number of streamlines with very similar anatomical neighbors. As a result, we could use a subset of these streamlines without affecting the computation of the anatomical priors. Therefore, for any WM bundle that included more than 20,000 training streamlines, we reduced that number to 20,000 to speed up this computation. We first removed outlier streamlines, which can be difficult to remove manually one by one, particularly for very dense bundles. Outliers were detected by mapping the end points of the streamlines to a common template space (see below for more information on registration), summing the endpoints over all subjects, and clustering them. Small clusters of endpoints were tagged as outliers and any individual streamlines that terminated in those outlier clusters were removed. If the total number of streamlines in a bundle was still above 20,000, it was reduced further by random subsampling of the streamlines. Note that this reduced set of streamlines was used to train TRACULA, but the complete set of 2.29 million streamlines was used as the “ground truth” to evaluate the accuracy of the automated reconstruction.

##### Anatomical neighborhood priors

For each subject, we used the 42 manually defined bundles from each of the other 15 subjects as the training set. The mathematical formulation has been described elsewhere (Yendiki et al. 2011; Yendiki et al. 2016). Briefly, this approach models a WM pathway as a cubic spline, which is initialized with the median streamline of the training set. A random sampling algorithm is used to draw samples from the posterior probability distribution of the pathway by perturbing the control points of the spline. The posterior probability is decomposed into the likelihood of the pathway given the DW volumes and the prior probability of the pathway. The likelihood term fits the shape of the spline to the diffusion orientations in the voxels that the spline goes through. As previously, diffusion orientations were obtained by fitting the ball-and-stick model (Behrens et al. 2003) to the subject’s DW volumes. This model does not require a sophisticated dMRI acquisition; it can be used on data collected with low b-values and with as few as 30 directions (Behrens et al. 2007).

The prior probability term in TRACULA fits the shape of a pathway to its anatomical neighborhood, given the manually labeled examples of this pathway from the training subjects and the anatomical segmentation volumes of both test and training subjects. Specifically, the training streamlines are used to compute the prior probability that each label of the anatomical segmentation is the *j*-th neighbor of the pathway at the *i*-th point along the trajectory of the pathway. Here *i* indexes equispaced points (3 mm apart) along the pathway and *j* indexes the nearest neighboring segmentation labels in different directions (left, right, anterior, posterior, etc.) The anatomical labels were extracted from the subject’s T1-weighted scan using FreeSurfer.

##### Structural segmentation

In this work, we used an anatomical segmentation volume that combined the labels of the Desikan-Killiany cortical parcellation (Desikan et al. 2006) with the standard FreeSurfer subcortical segmentation (Fischl et al. 2002). However, we replaced the thalamus label of the latter with the subject’s thalamic nuclei segmentation labels (Iglesias et al. 2015, 2018). This replacement was done to avoid oversegmenting the thalamus into WM voxels, and to provide additional specificity on the anatomical neighbors of tracts that terminate in or travel around the thalamus. Computing the prior probabilities on the anatomical neighbors of the tracts requires that each (training or test) subject’s anatomical segmentation be transformed to the subject’s individual dMRI space. This within-subject, dMRI-to-T1 alignment was performed by a boundary-based, affine registration method (Greve and Fischl 2009).

##### Template construction

Although finding the anatomical neighbors of a tract is a within-subject operation, it is important to ensure that all subjects’ brains have the same orientation, so that the relative positions of neighboring structures (which structure is to the left/anterior/etc. of which tract) is equivalent for all subjects. For this purpose, and for mapping the median of the training streamlines to the test subject during initialization, subjects must be mapped onto a template brain. Here we constructed a template by co-registering the FA maps of all 35 subjects in the MGH-USC HCP data set (Fan et al. 2015) with symmetric normalization (SyN; Avants et al. 2008), as implemented in ANTs (Avants et al. 2011). An affine initial registration was followed by 4 iterations of nonlinear registration with the b-spline SyN transform model, a cross-correlation similarity metric with a radius of 2, and a 4-level multi-resolution scheme with 100/70/50/50 sub-iterations per level. Each test subject’s FA map was aligned to the template with the default sequence of rigid/affine/deformable SyN registration followed in ANTs. Although we are introducing this nonlinear registration approach to TRACULA in the interest of generality, it is important to note that the purpose for which TRACULA performs subject-to-template registration (to find within-subject anatomical neighbors in a consistent set of directions) does not require exact voxel-wise, inter-subject alignment. We demonstrate this here by comparing this nonlinear registration approach to the one that was used by default in previous versions of TRACULA, *i.e.,* affine registration of each subject’s T1 image to the 1 *mm* MNI-152 template with FSL’s FLIRT (Jenkinson et al. 2002).

##### Choice of control points

The number of control points of the cubic spline, which are perturbed at each iteration of the random sampling algorithm to draw new sample paths, was chosen according to the average length of the training streamlines for each bundle. Specifically, we chose the number of control points to be 5 for the genu of the CC, and we then set the number of control points for all other bundles proportionally to their length. This ranged from 4 control points for the ATR to 12 control points for the temporal component of the body of the CC.

##### Along-tract analysis

Pointwise assessment of streamline tractography attributes (PASTA) is a type of analysis where an along-tract profile of a microstructural measure (*e.g.,* FA) is generated by averaging the values of the measure at different cross-sections of a tract (Jones et al. 2005). For each of the 42 bundles, we generated a reference streamline for PASTA analyses, to ensure that all subjects are sampled at the same number of cross-sections along a given bundle. The reference streamline was the mean of the manually annotated streamlines in template space. After the bundles of an individual subject were reconstructed automatically with TRACULA, the reference streamlines were mapped from the template to the individual. We generated along-tract profiles of microstructural measures by projecting the value of each measure from every point on every automatically reconstructed streamline to its nearest point on the reference streamline. Values projected to the same point on the reference streamline were then averaged, to generate an along-tract, 1D profile of the microstructural measure.

#### 2.5.2 Multi-ROI

For comparison, we also reconstructed each subject’s bundles with a commonly used multi-ROI approach, which maps a set of ROIs from a template to an individual subject’s dMRI space and combines them with a set of deterministic inclusion and exclusion rules to constrain the output of local probabilistic tractography (Groot et al. 2013; Warrington et al. 2020). For each subject, we used the ROIs that we had drawn for the manual labeling of the bundles in the other 15 subjects. We aligned the subjects to the FMRIB-58 FA template using FSL’s FNIRT, and then used the resulting nonlinear warp to transform the ROIs to template space. We summed the corresponding ROIs of the 15 subjects, and thresholded their sum to ensure that it had a size similar to that of the individual ROIs. (Empirically this was done by applying a lower threshold equal to 30% of the number of subjects). The group-averaged and thresholded ROIs were then mapped to the test subject using the inverse of the subject-to-template registration. For each pathway, the automated multi-ROI protocol used these ROIs as inclusion masks. For the bundles that were included in previously published multi-ROI protocols (Warrington et al. 2020), we used the previously proposed exclusion masks and augmented them as needed with the group-averaged exclusion masks from our own manual dissections. Local probabilistic tractography was performed using FSL’s probtrackX (Behrens et al. 2007) in symmetrical mode (seeding from both inclusion masks) with default parameters (5000 number of samples, 200 steps per sample, 0.5 *mm* step-length) and the same ball-and-stick model as in the previous section (Behrens et al. 2003). We implemented along-tract (PASTA) analyses for the multi-ROI approach, using the same reference streamlines as for TRACULA, in the manner described in section 2.5.1 above.

#### 2.5.3 Accuracy of automated reconstruction

We assessed the accuracy of the TRACULA and multi-ROI automated reconstruction by comparing the tracts reconstructed automatically in the b=1000 *s*/*mm*^2^, 64-direction data to those labeled manually in the b_max_=10,000 *s*/*mm*^2^, 512-direction data of the same subject. We quantified the reconstruction error by computing the modified Hausdorff distance (MHD; Dubuisson and Jain, 1994) between the automatically reconstructed and manually labeled pathways. The MHD between two set of points *S* and *T* is defined as the minimum distance between a point in one set and any point in the other set, averaged over all points in the two sets:

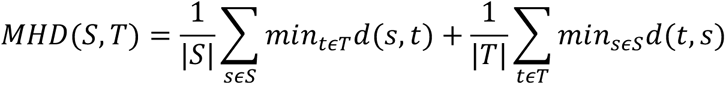

where *d*(·,·)is the Euclidean distance between a pair points from the two sets, and |·| is the size of a set. Greater MHD indicates greater deviation of the automatically reconstructed tract from the one labeled manually in the same subject, and hence lower accuracy of the automated reconstruction.

In previous work, we reported MHD of tracts reconstructed with TRACULA using our older training sets for adult brains (Yendiki et al. 2011) or infant brains (Zöllei et al. 2019), after thresholding the voxel visitation maps of the automatically reconstructed tracts at a single threshold (20% of the maximum, which is the default visualization threshold in TRACULA). However, for the purpose of a comparison between TRACULA and the multi-ROI approach, a single threshold would not be informative. The global tractography used in TRACULA adds an entire end-to-end path to the voxel visitation map at each iteration, whereas the local tractography used in the multi-ROI approach adds a single voxel at every iteration. As a result, thresholding at the same percentage of the peak value does not yield equivalent results between the two methods. For this reason, in the experiments presented here we performed a more comprehensive evaluation of reconstruction error, where we increased the threshold gradually from 0% to 90% for both methods, and computed their MHD at each threshold.

In addition, for each bundle and at each threshold, we computed the true-positive rate (TPR), which quantifies the proportion of the manually labeled streamlines that overlap with the automatically reconstructed bundle:

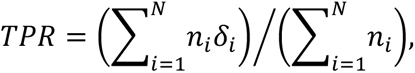

where *n_i_* the number of manually labeled streamlines that go through the *i*-th voxel, *δ_i_* an indicator function that is equal to 1 if the automatically reconstructed bundle goes through the *i*-th voxel and 0 otherwise, and *N* the number of voxels in a brain volume. Each true positive voxel (*δ_i_* = 1) is weighed by the number of manually labeled streamlines *n_i_* that go through that voxel, to account for the fact that the manually labeled bundles themselves contain noisy tractography streamlines. Thus, a true positive should be rewarded more if it occurs in a voxel that overlaps with a large number of the manually labeled streamlines.

In a conventional receiver operating characteristic (ROC) analysis, the TPR is plotted against the false-positive rate (FPR), which quantifies the proportion of the automatically reconstructed bundle that does not overlap with the manually labeled one:

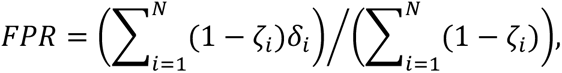

where *ζ_i_* an indicator function that is equal to 1 if the manually labeled bundle goes through the *i*-th voxel and 0 otherwise. It is important to note, however, that the FPR penalizes all false positive voxels equally, no matter how far away from the manually labeled bundle they occur. Thus the MHD, which measures the distance between the automatically reconstructed and manually labeled bundles, is a more informative metric of reconstruction errors.

The goal of these experiments was to investigate how close automated tractography in routine-quality data could come to manually annotated tractography in high-quality data, hence the “ground truth” was obtained from the manually labeled, multi-ROI tractography of section *2.4*. However, there were cases where even the full b_max_=10,000 *s*/*mm*^2^ data yielded only a few streamlines for a certain manually labeled bundle. In those cases, measuring the accuracy of the automated reconstructions by comparison to the manually labeled bundle could underestimate the accuracy of the automated reconstruction. We identified such cases as manually labeled bundles whose volume was less than 1/3 of the median volume of the same bundle across the 16 training subjects. They were one case each of the LH-CBD, LH-AR, and CC-BODY-T, and two cases of the AC. We excluded these cases when computing the metrics described above, but including them would not change any of our conclusions.

#### 2.5.4 Test-retest reliability of automated reconstruction

We divided the 64 diffusion directions of the b=1000 *s*/*mm*^2^ shell into two subsets, each containing 32 directions that were approximately uniformly distributed over the sphere. We applied the automated reconstruction methods described in 2.5.1 and 2.5.2 to each of the subsets, and we computed the accuracy metrics of 2.5.3. This allowed us to assess if the results from the two methods were reproducible between the test and retest scans, and how robust the methods were to even lower angular resolution.

#### 2.5.5 Test-retest reliability of along-tract measures

For the bundles reconstructed from each of the two 32-direction datasets, either with TRACULA or with the multi-ROI method, we extracted PASTA profiles of FA and mean/radial/axial diffusivity (MD/RD/AD). We assessed the test-retest reliability of these profiles by computing the symmetrized percent change (SPC) between the profiles obtained by the same method from the two 32-direction datasets:

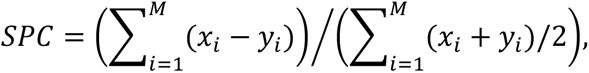

where *x_i_* and *y_i_* the *i*-th along-tract data point of a microstructural measure (FA/MD/RD/AD) from the two 32-direction datasets. The total number of data points, *M*, equals the number of cross-sections along a tract times the number of subjects.

We computed the test-retest reliability, as quantified by SPC, at a fixed level of sensitivity for both reconstruction methods. For the multi-ROI method, we set the threshold for the voxel visitation maps to 1% of the maximum value. At that threshold, the multi-ROI method had a sensitivity of about 0.6. We then set the threshold for TRACULA (10%) to achieve the same sensitivity.

#### 2.5.6 Evaluation on a larger dataset

As a final evaluation, we show preliminary results from assessing the ability of TRACULA to detect subtle microstructural effects in a larger dataset. We used data from 204 adolescents scanned for the Boston Adolescent Neuroimaging of Depression and Anxiety (BANDA) study, a Connectomes Related to Human Disease (CRHD) project. The study cohort had been recruited to probe the full continuum of depressed and anxious symptoms and their co-morbidity, and thus allow transdiagnostic investigations of brain-behavior relationships. It included 138 participants with depression and/or anxiety disorders (age 15.50±0.83, 95 female) and 66 controls (age 15.17±0.83 years, 36 female). Details on the clinical assessment and imaging protocol are provided elsewhere (Hubbard et al. 2020; Siless et al. 2020).

Here we used the T1-weighted images (. 8 *mm* isotropic resolution) to obtain structural segmentations with FreeSurfer; and the lower shell of the dMRI data (1.5 *mm* isotropic resolution, b=1500 *s*/*mm*^2^, 93 diffusion weighted volumes collected with two phase-encode directions each, and 28 non-diffusion weighted volumes) to reconstruct WM pathways with TRACULA. The dMRI data were pre-processed with FSL’s *topup* (Andersson et al. 2003) and *eddy* (Andersson & Sotiropoulos 2016) to mitigate susceptibility and eddy-current distortions. We reconstructed the following pathways with TRACULA: all subdivisions of the CC, and bilateral ATR, CBD, CBV, EmC, FX, SLF1, SLF2, SLF3, UF. We studied these pathways as they have been previously reported to be affected in patients with depression or anxiety (Bracht et al., 2015; Greenberg et al., 2021; Henderson et al., 2013; LeWinn et al., 2014; Liao et al., 2014). We tested the along-tract FA values for associations with three clinical variables: the total score from the Mood and Feelings Questionnaire (MFQ; Angold et al. 1995) and the depression and general anxiety subscale scores from the Revised Child Anxiety and Depression Scale (RCADS; de Ross et al. 2000). We excluded two participants out of the full cohort of 206 due to missing clinical scores.

For each clinical score, we fit a general linear model (GLM) with the along-tract FA value as the dependent variable, and sex, age, and clinical score as the independent variables. We tested two contrasts for statistical significance: the average slope of FA vs. clinical score, and the difference of slopes between female and male participants. We used FreeSurfer statistical analysis tools, adapted for 1D data; specifically, we fit a GLM at each point along each tract with *mri_glmfit*, and performed simulation-based, cluster-wise correction for multiple comparisons with *mri_glmfit-sim* (Hagler et al. 2006; Greve and Fischl 2018). The cluster-forming threshold and the cluster-wise threshold for statistical significance were both set to *p*=0.05, and 1000 simulations were performed. After statistical testing, we visualized the along-tract *p*-values by projecting them onto a randomly selected subset of the training streamlines in template space.

## 3. Results

### 3.1 Manually labeled dataset

Fig. 5 shows the 42 manually labeled pathways. The full set includes 2.29 million annotated streamlines. In individual dMRI space, they cover 82% of all cerebral and cerebellar WM voxels across the 16 subjects. For Fig. 5, the streamlines were mapped to template space and aggregated across all 16 training subjects. In template space, 98% of cerebral and cerebellar WM voxels (defined by majority voting of the anatomical segmentations of the 16 subjects) overlap with the streamlines of at least one subject. Thus, although the 42 pathways that we have labeled here do not represent all brain connections, they provide extensive WM coverage.

**Fig. 5.**
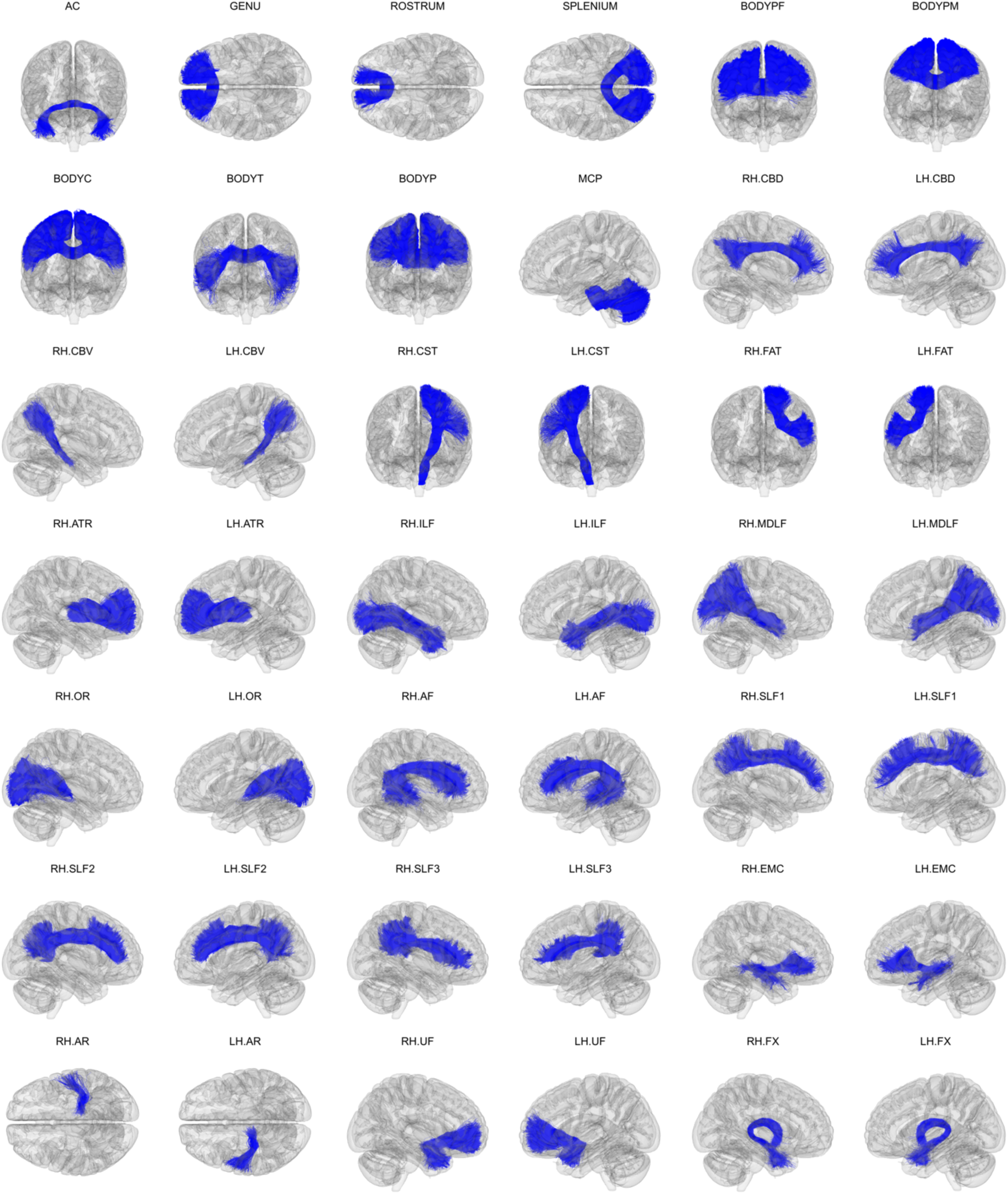
Manually labeled dataset. Manually labeled streamlines from each of the 42 WM bundles are shown aggregated over all 16 training subjects. Manual annotation was performed on each subject’s individual dMRI data as described in section 2.4. Streamlines are displayed here in 1 *mm* MNI-152 template space.

Fig. 6 shows the coverage of the cortical surface by the terminations of the manually labeled streamlines. For this figure, the number of streamline end points per voxel were summed along the normal of the surface, within 3mm from the WM-GM junction. They were then mapped from each individual’s surface to the *fsaverage* surface using the FreeSurfer spherical morph. The total numbers of streamlines across the 16 subjects were then obtained at each vertex. No smoothing was applied in the volume or on the surface to produce these maps. The terminations of the manually labeled streamlines cover 89% of the cortical surface on the left hemisphere and 88% on the right hemisphere.

**Fig. 6.**
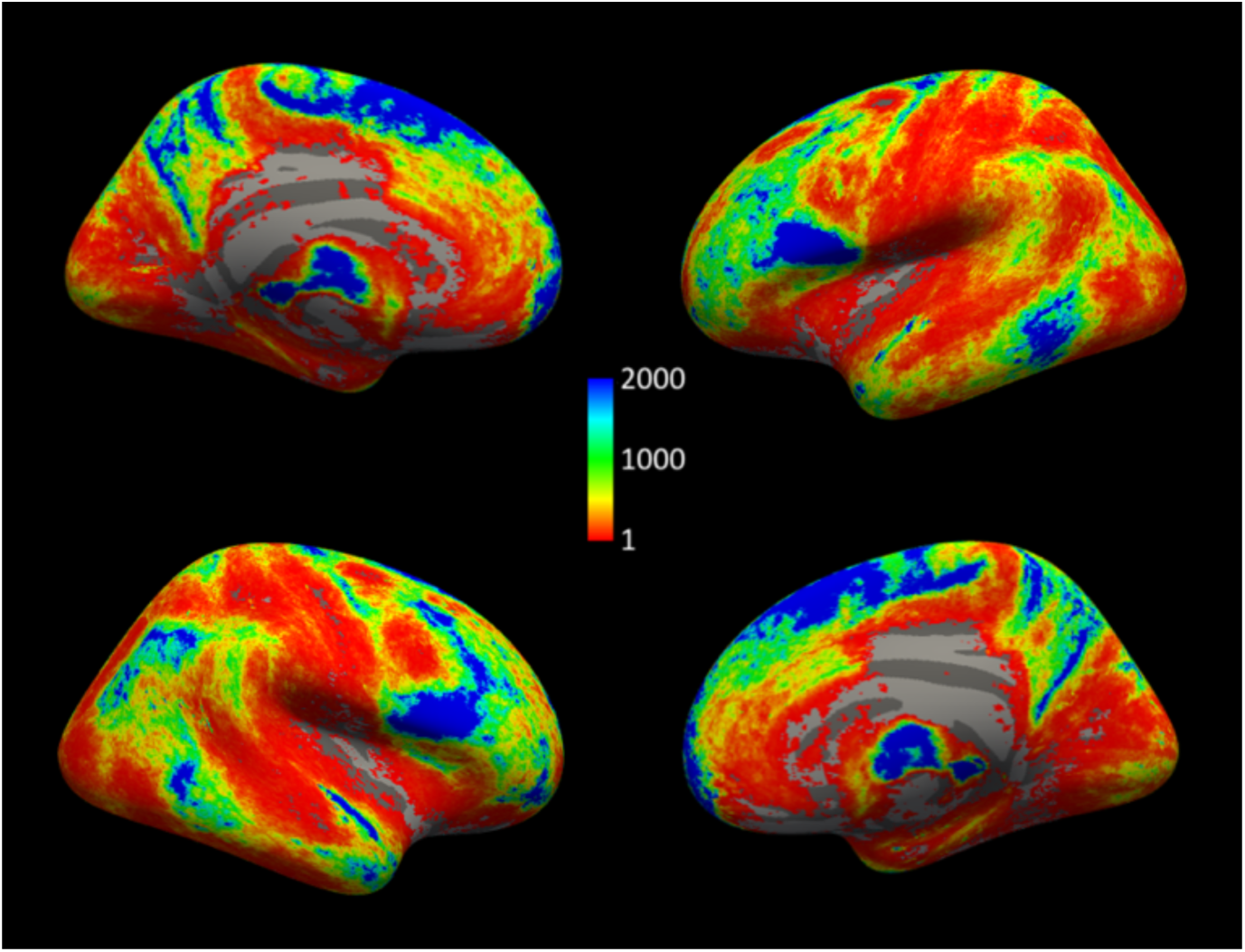
Cortical terminations of manually labeled streamlines. Total number of the streamlines in the manually labeled set that terminate within 3 *mm* of each vertex on the WM-GM boundary in fsaverage space.

Fig. 7 shows the FA template that we constructed from the 35 MGH-USC HCP subjects and that we used as the target for inter-subject registration with ANTs. The figure also shows the mean of the manually annotated streamlines from each of the 42 WM bundles. We used these mean streamlines as the reference streamlines for PASTA analysis.

**Fig. 7.**
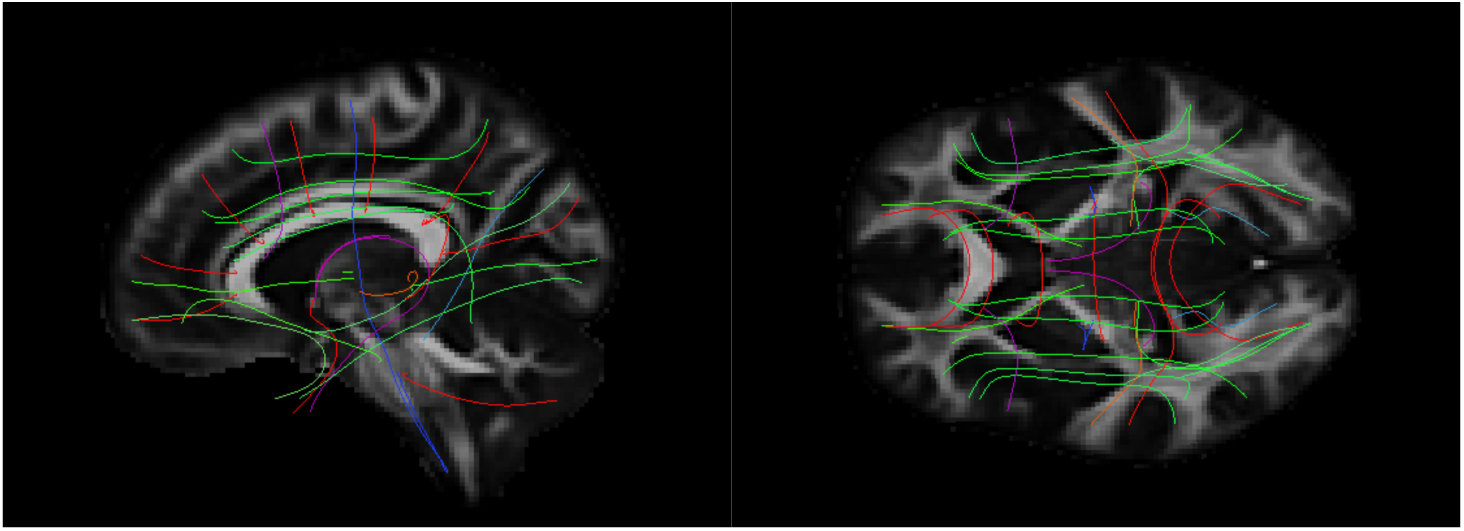
Template and reference streamlines. The template that we constructed from the FA maps of all 35 MGH-USC subjects is shown in sagittal (left) and axial (right) view. The mean of the manually annotated streamlines from each of the 42 bundles is also shown. These serve as the reference streamlines where microstructural measures are projected for PASTA analyses.

### 3.2 Comparison of automatically reconstructed and manually labeled pathways

Fig. 8 shows the accuracy measures of section 2.5.3, computed over all 42 pathways and 16 subjects in the leave-one-out experiments. Results are shown for the 64-direction, b=1000 *s*/*mm*^2^ data with TRACULA (red) and the multi-ROI method (black); and for two sets of 32-direction, b=1000 *s*/*mm*^2^ data with TRACULA (yellow, green) and the multi-ROI method (blue, purple). The plot on the left shows the sensitivity (TPR) as a function of 1-specificity (FPR). The plot on the right shows the reconstruction error (MHD in mm) as a function of sensitivity. Mean MHD is shown with standard error bars. Each point along the curves represents a different threshold applied to the probability distributions estimated by each method. The point of highest sensitivity is the one achieved by unthresholded distributions.

**Fig. 8.**
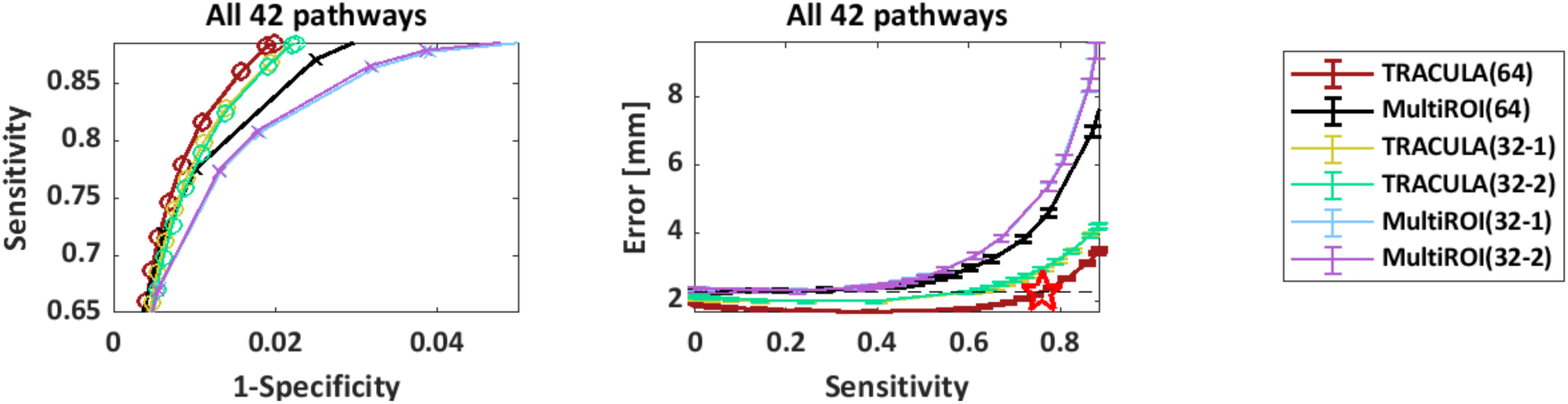
Overall accuracy of automated reconstruction. For each reconstruction method (TRACULA, multi-ROI), results are shown for 64 directions and for 2 sets of 32 directions. Measures were computed across all 42 pathways and 16 manually labeled subjects. Each point along the curves represents a different threshold applied to the estimated probability distributions. **Left:** Sensitivity (TPR) vs. 1-specificity (FPR). **Right:** Reconstruction error (MHD in mm) vs. sensitivity. Horizontal dashed line: minimum MHD achieved by the multi-ROI method on the 64-direction data. Red star: Maximum TPR achieved by TRACULA at the same MHD level.

The highest sensitivity achieved by TRACULA across all 42 pathways was 89%, indicating high coverage of the “ground-truth” pathways, *i.e.,* the ones obtained from the manual labeling of the 512-direction, b_max_=10,000 *s*/*mm*^2^ data. At that sensitivity, the reconstruction error (MHD) was 3.5 mm for TRACULA on the 64-direction data. Compared to that, the reconstruction error at the same sensitivity level was 4.2 mm (20% higher) for TRACULA on both sets of 32-direction data, 7.6 mm (118% higher) for the multi-ROI method on the 64-direction data, and 10.6/10.4 mm (203/197% higher) for the multi-ROI method on the two sets of 32-direction data. For both reconstruction methods, the overall performance metrics were highly reproducible between the two sets of 32-direction data. This is illustrated by the overlap of the green and yellow curves (for TRACULA) and the overlap of the blue and purple curves (for the multi-ROI method). For both methods, performance was somewhat lower on the 32-direction data than the 64-direction data. The multi-ROI method exhibited a greater deterioration as a result of decreasing the number of directions from 64 to 32.

Figs. 9-11 show plots of the reconstruction error (MHD) vs. sensitivity (TPR) separately for each of the 42 pathways. There was some variability across pathways in terms of the difference in performance between reconstruction methods, the extent to which lowering the number of directions from 64 to 32 affected their performance, or the level of reproducibility between the two sets of 32 directions. However, the general patterns observed from the overall performance plot of Fig. 8 could also be observed from the individual pathway plots of Figs. 9-11.

**Fig. 9.**
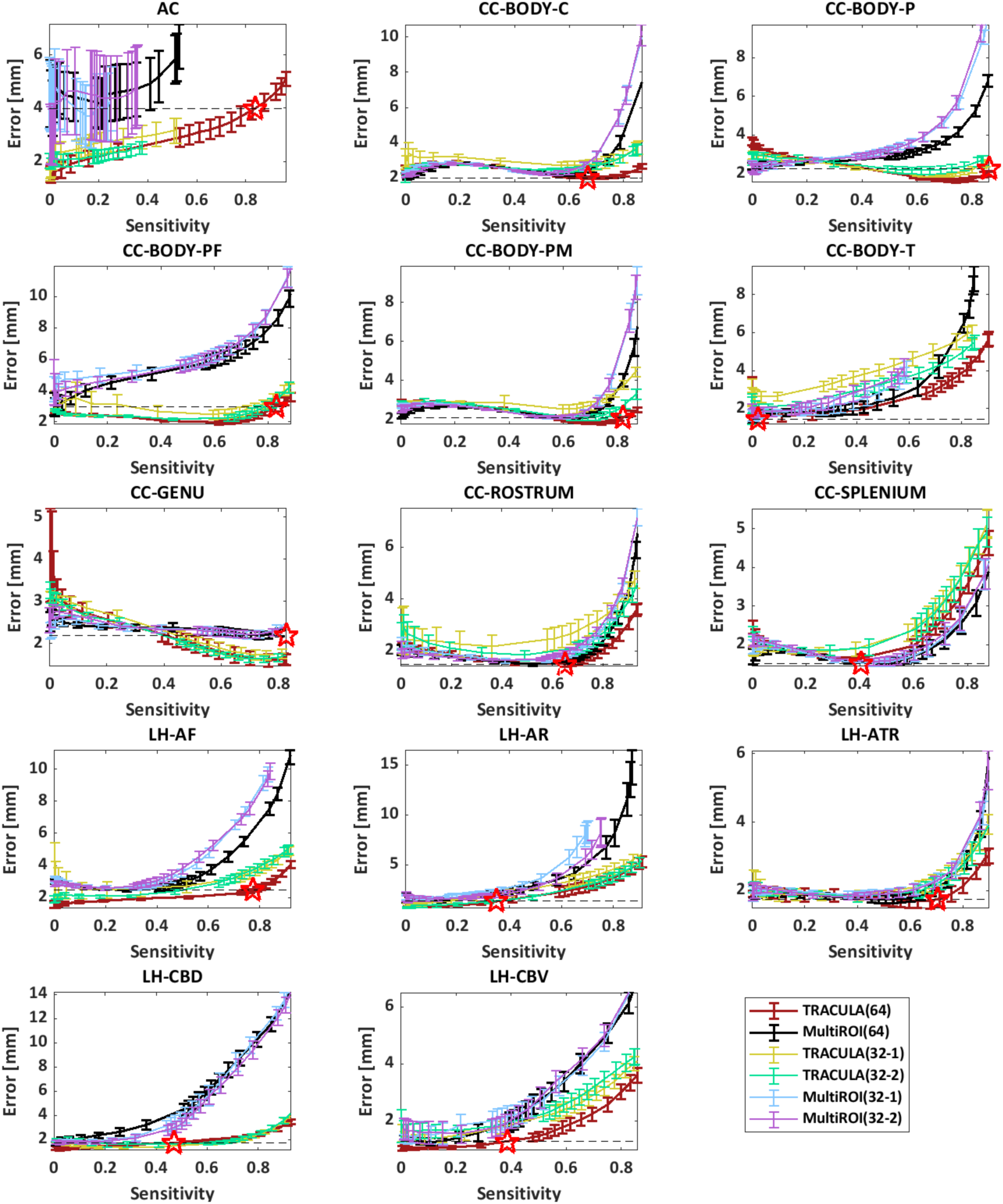
Accuracy of automated reconstruction by pathway. For each reconstruction method (TRACULA, multi-ROI), results are shown for 64 directions and for 2 sets of 32 directions. Each point along the curves represents a different threshold applied to the estimated probability distributions. **Left:** Sensitivity (TPR) vs. 1-specificity (FPR). **Right:** Reconstruction error (MHD in mm) vs. sensitivity. Horizontal dashed line: minimum MHD achieved by the multi-ROI method on the 64-direction data. Red star: Maximum TPR achieved by TRACULA at the same MHD level.

**Fig. 10.**
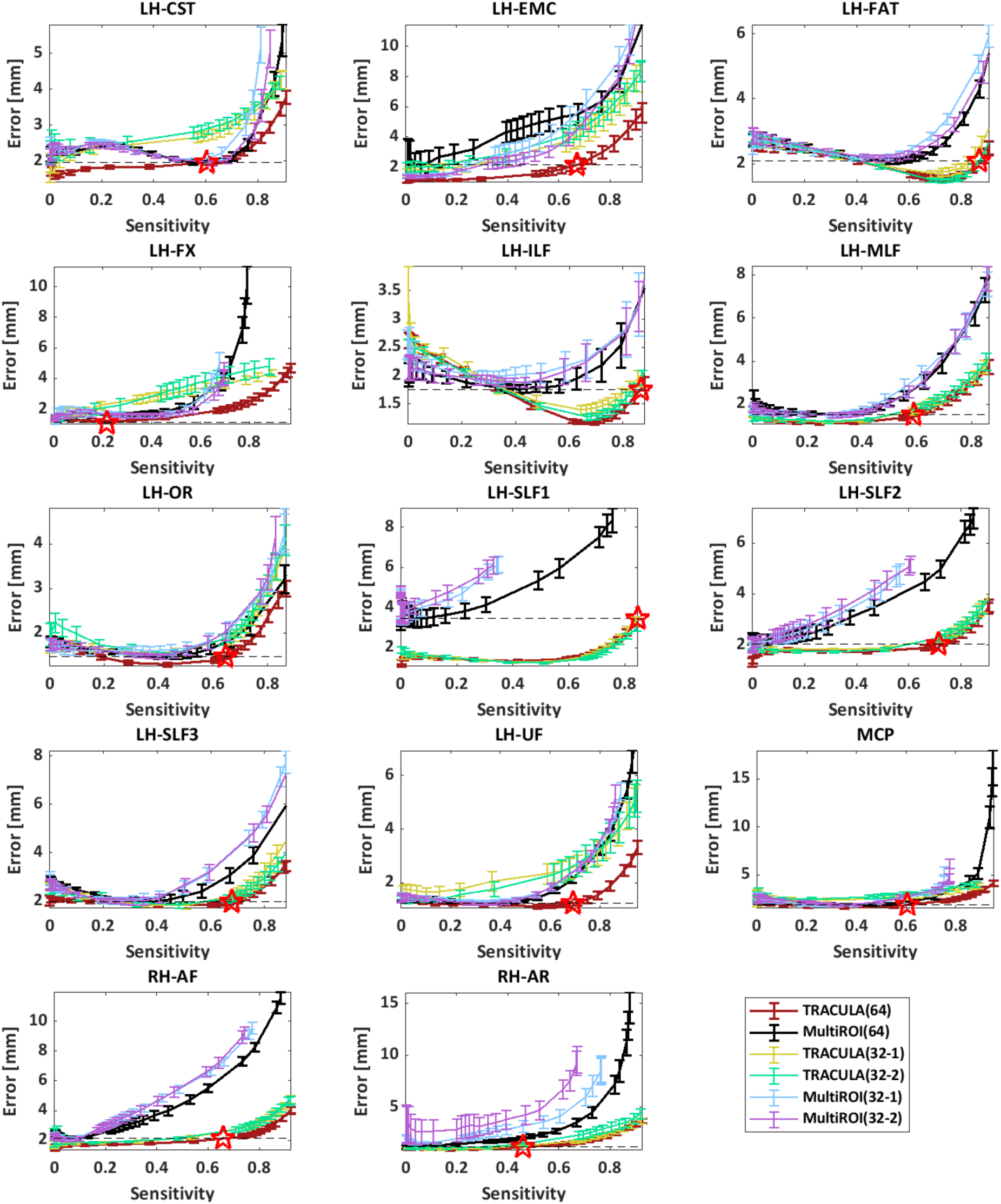
Accuracy of automated reconstruction by pathway (continued). For each reconstruction method (TRACULA, multi-ROI), results are shown for 64 directions and for 2 sets of 32 directions. Each point along the curves represents a different threshold applied to the estimated probability distributions. **Left:** Sensitivity (TPR) vs. 1-specificity (FPR). **Right:** Reconstruction error (MHD in mm) vs. sensitivity. Horizontal dashed line: minimum MHD achieved by the multi-ROI method on the 64-direction data. Red star: Maximum TPR achieved by TRACULA at the same MHD level.

**Fig. 11.**
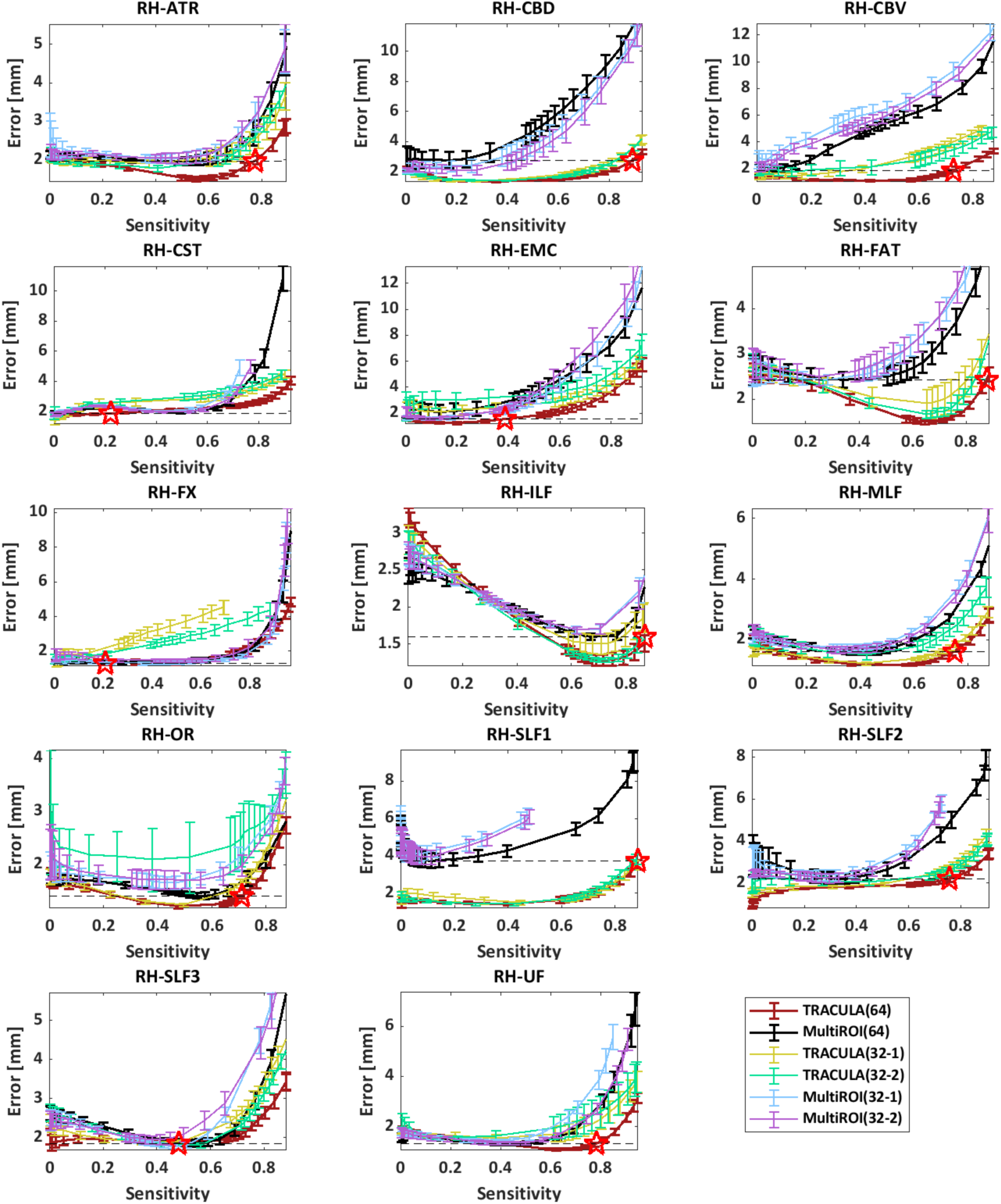
Accuracy of automated reconstruction by pathway (continued). For each reconstruction method (TRACULA, multi-ROI), results are shown for 64 directions and for 2 sets of 32 directions. Each point along the curves represents a different threshold applied to the estimated probability distributions. **Left:** Sensitivity (TPR) vs. 1-specificity (FPR). **Right:** Reconstruction error (MHD in mm) vs. sensitivity. Horizontal dashed line: minimum MHD achieved by the multi-ROI method on the 64-direction data. Red star: Maximum TPR achieved by TRACULA at the same MHD level.

In the plots of reconstruction error (MHD) vs. sensitivity (TPR) from Figs. 8-11, a horizontal dashed line indicates the minimum MHD that can be achieved by the multi-ROI method on the 64-direction data, *i.e.,* the minimum MHD along the black curve. The portion of the red curve that lies below the dashed line represents the range of operating points for which TRACULA achieved a reconstruction error equal or less than the minimum achieved by the multi-ROI method. The red star indicates the maximum sensitivity that TRACULA could achieve while staying below that level of reconstruction error. Fig. 12 shows the sensitivity (TPR) values at these operating points. The gray bars show the sensitivity of the multi-ROI method at the threshold where it achieves its minimum reconstruction error. The red bars show the maximum sensitivity that TRACULA could achieve while maintaining a reconstruction error equal or less than the minimum error achieved by the multi-ROI method (*i.e.,* the sensitivity of TRACULA at the points marked by red stars in Figs. 8-11).

**Fig. 12.**
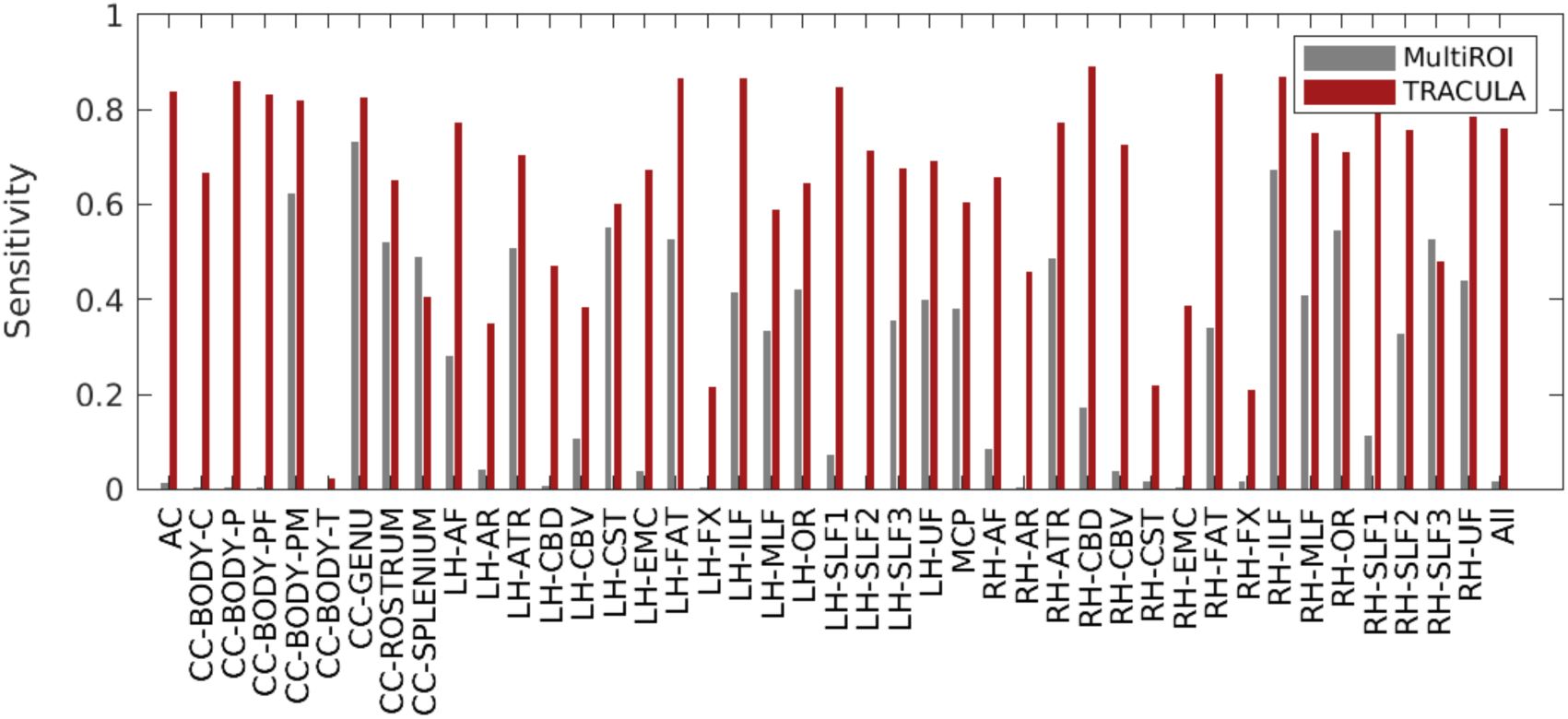
Maximum sensitivity at the same level of reconstruction error. For each of 42 pathways (and across all pathways on the far right), the plot shows the sensitivity (TPR) that the multi-ROI method achieves when its threshold is chosen to minimize the reconstruction error (MHD), and the maximum sensitivity that TRACULA can achieve while maintaining the same or lower reconstruction error.

Fig. 13 shows the minimum reconstruction error, as quantified by the MHD in mm, achieved by the multi-ROI method and TRACULA for each pathway. The x=y line is shown in black dots. The data points fall mostly above the x=y line, indicating that the minimum error was smaller for TRACULA than the multi-ROI method. Note that these errors do not correspond to matched thresholds or matched sensitivity levels between the two methods. They are the minimum errors that each method could achieve across all thresholds and thus sensitivity levels. Figs. 8-11 show that, when compared at matched levels of sensitivity, TRACULA could achieve overall lower reconstruction errors.

**Fig. 13.**
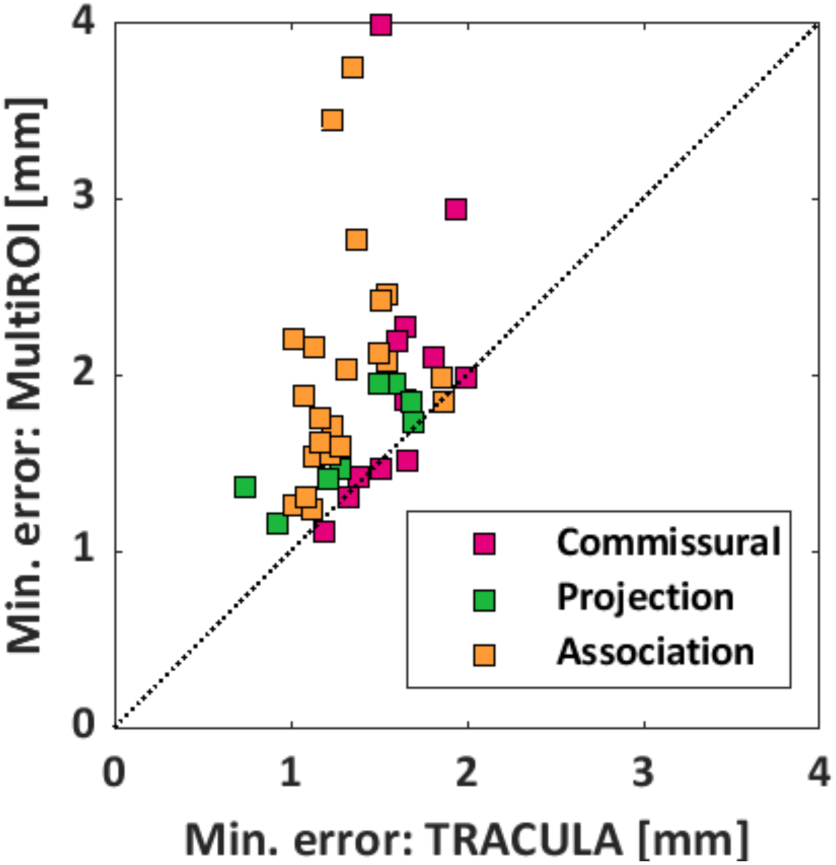
Minimum reconstruction error. For each of 42 pathways, the plot shows the minimum reconstruction error (MHD in mm) that can be achieved by TRACULA (x-axis) and the multi-ROI method (y-axis). The pathways are color-coded based on their type (commissural, projection, or association).

Fig. 14 shows that the performance of TRACULA is independent of the method that it uses for inter-subject registration. The plots show results from automated reconstruction on the 64-direction data with three methods: TRACULA or the multi-ROI method with nonlinear inter-subject registration (same as in Fig. 8), and TRACULA with affine inter-subject registration. As seen in the plots, performance is indistinguishable between TRACULA with the two registration approaches. This is because the anatomical priors in TRACULA do not encode information about the absolute coordinates of the pathways in template space. They only encode information about the relative positions (left, right, anterior, *etc.*) of the pathways with respect to their surrounding anatomical structures.

**Fig. 14.**
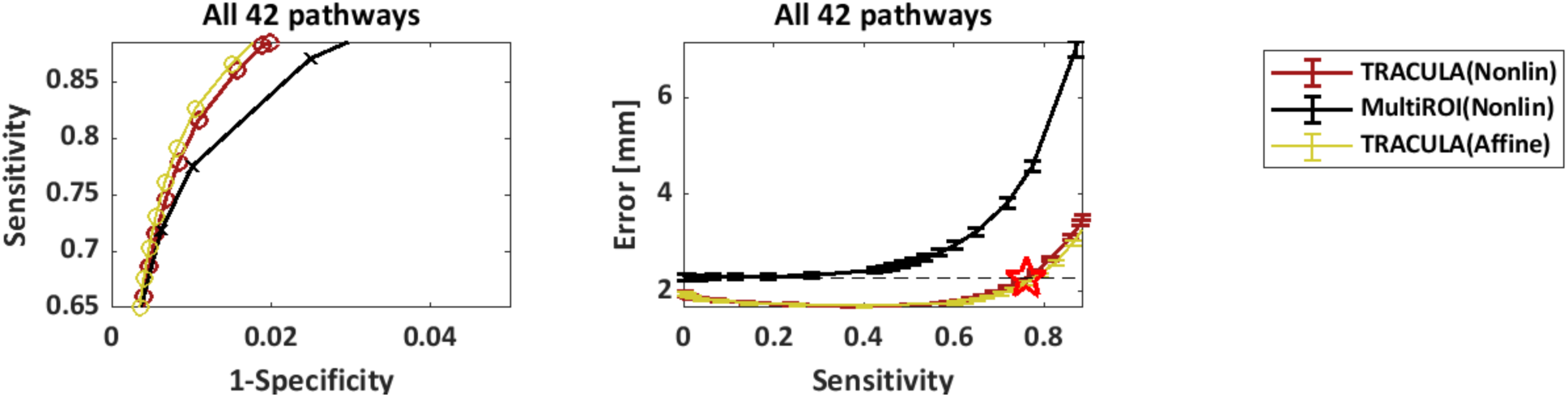
Robustness to inter-subject registration. Results are shown for reconstruction on the 64-direction data using either TRACULA or the multi-ROI method with nonlinear inter-subject registration (same as in Fig. 8), as well as TRACULA with affine inter-subject registration. Measures were computed across all 42 pathways and 16 manually labeled subjects. Each point along the curves represents a different threshold applied to the estimated probability distributions. **Left:** Sensitivity (TPR) vs. 1-specificity (FPR). **Right:** Reconstruction error (MHD in mm) vs. sensitivity. Horizontal dashed line: minimum MHD achieved by the multi-ROI method on the 64-direction data. Red star: Maximum TPR achieved by TRACULA at the same MHD level.

### 3.3 Test-retest reliability of along-tract measures

Fig. 15 shows the SPC of along-tract FA values between the two 32-direction datasets, for TRACULA and the multi-ROI method, at a sensitivity level of 0.6. An analysis of variance with factors of bundle (42 levels) and reconstruction method (2 levels) showed a significant effect of both bundle (*p*=2.9e-04) and reconstruction method (*p*=3.9e-08). Very similar results were obtained for MD (bundle: *p*=4.3e-03; reconstruction method: *p*=6.7e-08), RD (bundle: *p*=4.3e-03; reconstruction method: *p*=5.1e-08), and AD (bundle: *p*=7.8e-03; reconstruction method: *p*=5.4e-08).

**Fig. 15.**
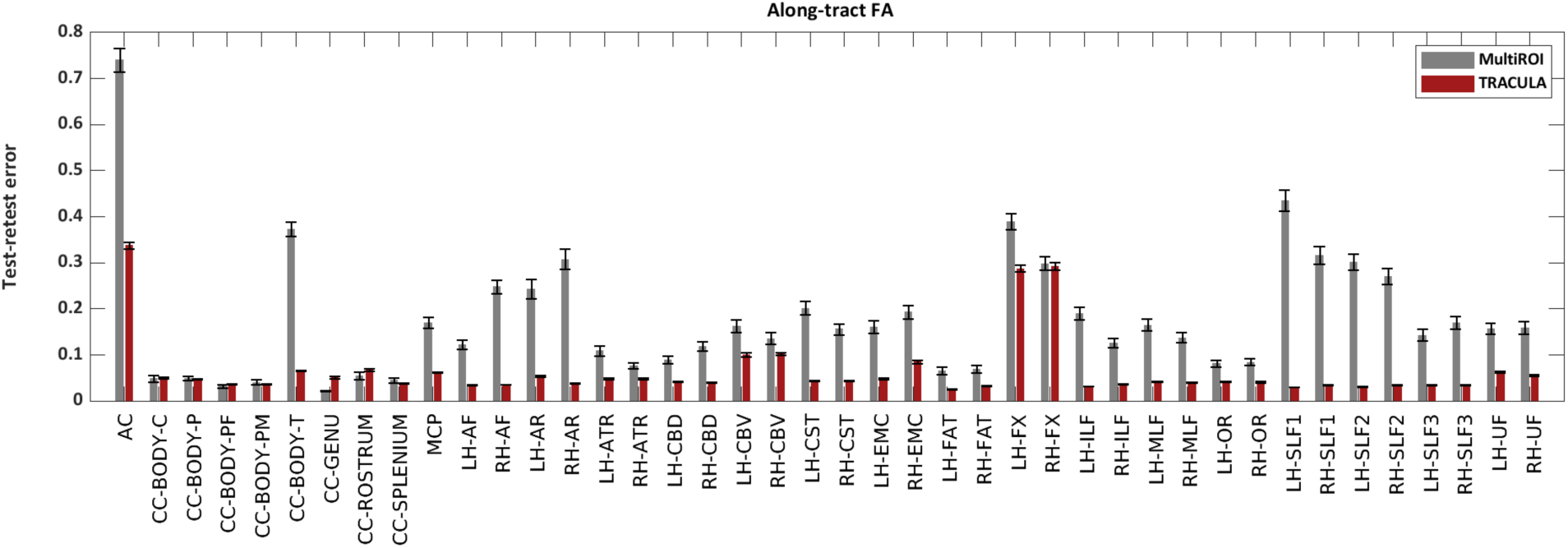
Test-retest reliability of along-tract FA. The plots show the test-retest error of along-tract (PASTA) FA values, as quantified by the SPC between along-tract FA obtained from two 32-direction data sets, with the multi-ROI method (gray) or with TRACULA (red). For both methods, pathway probability maps were thresholded to achieve a sensitivity of 0.6.

These results reflect both the reliability of automated tractography and the reliability of the microstructural measures themselves. For example, the two bundles where along-tract FA/MD/RD/AD had their lowest reliability (AC and FX) were the ones where these tensor-based measures would be the most prone to partial voluming due to proximity to CSF. Microstructural measures extracted from models other the tensor may be more reliable than these overall. Here, however, our main interest was in the comparison of reliability between the two reconstruction methods. The median test-retest error across all 42 bundles was 4.3% (FA), 2.6% (MD), 5.7% (RD), 3.2% (AD) for TRACULA; and 15.7% (FA), 12.4% (MD), 17.0% (RD), 12.9% (AD) for the multi-ROI method.

### 3.4 Evaluation on a larger dataset

Fig. 16 shows findings from the statistical analysis of along-tract FA in the 204 subjects of the BANDA cohort. The top row shows the WM bundles where the average slope of along-tract FA vs. clinical score was statistically significant. We found a negative slope of FA vs. clinical score in the LH-SLF1, for all three clinical scores (MFQ, RCADS-Dep, RCADS-GenAnx). The bottom row shows the bundles where the difference in slopes between female and male participants was statistically significant. We found greater slopes in females than males for MFQ vs. FA in the CC-BODY-PM; RCADS-Dep vs. FA in the CC-BODY-PM, RH-SLF1; and RCADS-GenAnx vs. FA in the RH-EMC, RH-FX, RH-CBD, LH-CBD, RH-ATR.

**Fig. 16.**
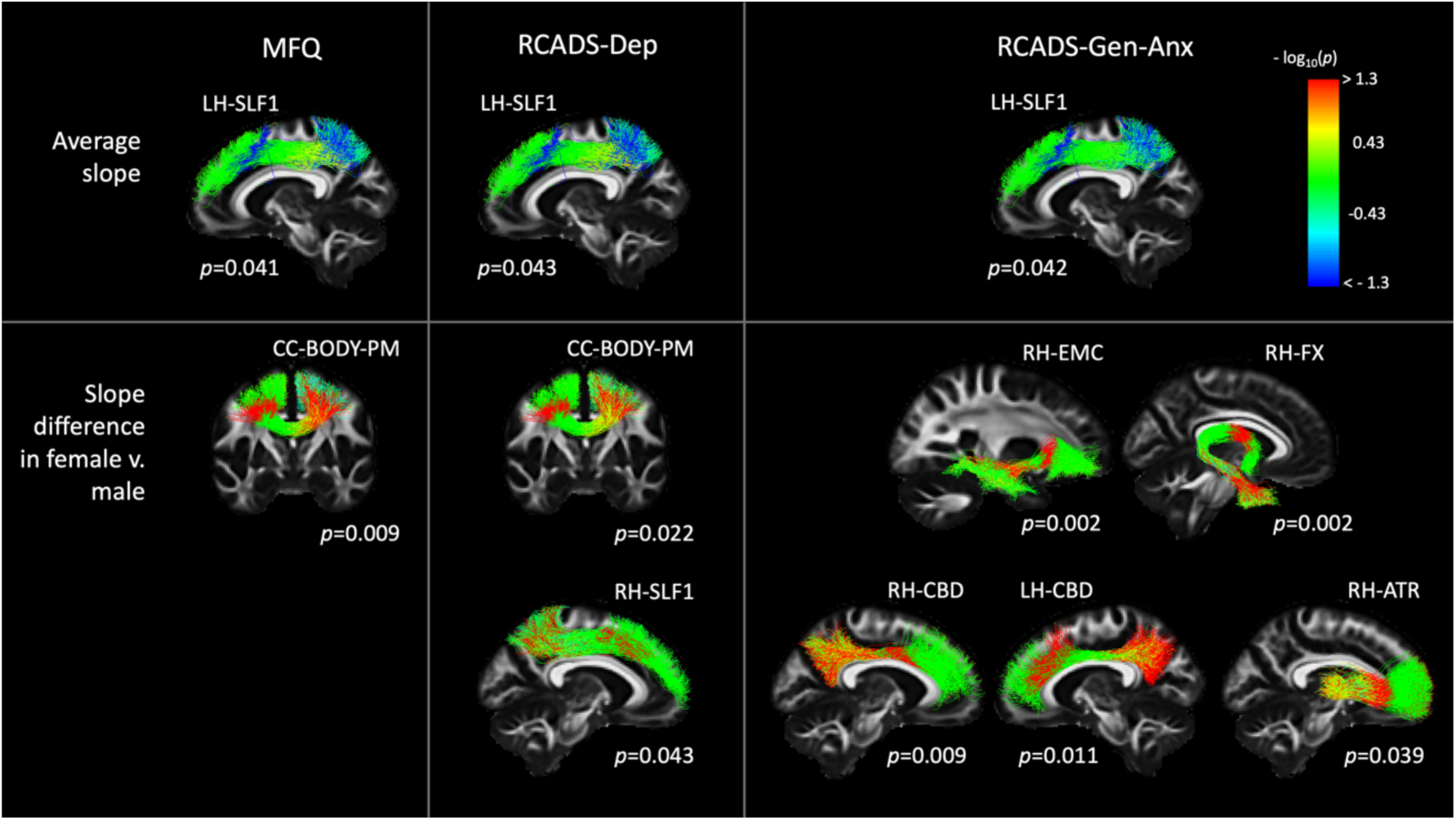
Associations of along-tract FA with clinical scores in the BANDA cohort. Each column shows results from a different clinical score (MFQ, RCADS-Dep, RCADS-Gen-Anx). Each row shows results from a different contrast (top: average slope of FA vs. clinical score; bottom: difference in slopes of FA vs. clinical scores between female and male participants). Pathways were reconstructed automatically with TRACULA. For display, along-tract p-values were mapped onto a randomly selected subset of the training streamlines in template space.

## 4. Discussion

In this work we present a new set of protocols for manual labeling of 42 major WM pathways using probabilistic tractography on high-quality (b_max_=10,000 *s*/*mm*^2^, 512-direction) dMRI data from a Connectom scanner. We also demonstrate that these manually annotated pathways can be used as training data to reconstruct the same pathways automatically from routine-quality (b=1000 *s*/*mm*^2^, 64-direction) dMRI data with high sensitivity and high reliability.

### 4.1 Manual labeling

The widely used protocols for manual labeling of WM pathways were introduced at a time when tractograms were typically obtained by running deterministic tensor tractography on dMRI data with low b-values and low angular resolution (Wakana et al. 2007; Catani and Thiebaut de Schotten 2008). These protocols were a critical step towards applying dMRI tractography to population studies. They introduced the concept of the multi-ROI tract dissection, which was also the first method used for automated tract-of-interest reconstruction (W. Zhang et al. 2008; Clayden et al. 2009).

Since then, the acquisition technologies adopted and advanced by the HCP led to a dramatic improvement in the quality of *in vivo* dMRI data. The higher spatial and angular resolution of modern dMRI data, coupled with the use of probabilistic tractography and crossing-fiber modeling techniques, yield much larger and more complex tractograms. These can be used for a more detailed and accurate definition of WM pathways, but they also contain many more noisy streamlines and require more clean-up. While the previously proposed manual annotation protocols are an excellent starting point, they need to be updated with a greater number of inclusion and exclusion ROIs. Furthermore, some pathways that were not typically included in older “virtual dissection” protocols, because they could not be reconstructed reliably with older data, can now be readily extracted from modern tractograms.

In section 2.4, we presented an updated set of protocols that we deployed to label 42 WM pathways manually in the MGH-USC HCP data. These data, which could only be acquired with a Connectom scanner, allowed a more detailed and accurate reconstruction of major brain pathways, as they had been described in anatomical studies. We were able to obtain a more comprehensive delineation of the termination regions of these pathways, and to reconstruct bundles or portions of bundles that were not accessible before, like the acoustic radiation (Maffei et al. 2019), the more lateral terminations of the CST in the motor cortex, or the Meyer loop of the OR.

However, our ability to reconstruct certain aspects of the more challenging WM bundles is still limited, even with the best available in vivo dMRI data. Here we discuss some examples of discrepancies between tractography on high-quality dMRI data and the anatomical literature because we believe that they can be useful benchmarks for developers of tractography algorithms and useful targets for future investigation with ex vivo dMRI. These examples are from the AC, ATR, CST, FX, UF, and SLFI.

#### AC

The AC is a thin, long compact bundle with an uncommon C-shape that connects the two temporal lobes (J. Schmahmann and Pandya 2006). In its course, the AC lies in the proximity of the putamen, caudate nucleus, globus pallidus, amygdaloidal nuclei, and temporal and perirhinal cortex. The vicinity to these GM structures makes the AC sensitive to partial volume effects, which can severely affect its reconstruction, especially given its small size (only a few voxels wide). In the temporal lobe, the AC fibers fan out towards the anterior part of the temporal pole, where they merge with the fibers of the UF and FX (Cavdar et al. 2020). This configuration, in which different fiber bundles merge and intermingle, is hard to resolve with tractography, and it usually results in favoring the reconstruction of the bigger bundles that intersect with the AC. While we could reconstruct the AC correctly in most of the 16 subjects, some presented only a few valid streamlines, and in most subjects the temporal terminations were sparse and noisy.

#### ATR

We defined the ATR as cortico-thalamic fibers connecting the thalamus to the frontal cortex. We recognize that this definition remains vague and reflects a tractography-based characterization of this bundle more than an anatomical one (Safadi et al., 2018). Because of the limitations of diffusion tractography, we are not able to precisely separate these fibers from the fibers projecting from/to the brainstem, and we therefore recognize the possibility that some of the latter fibers are also included in the delineation of the ATR. We also observed that in all our manual dissections it was difficult to obtain the most dorsolateral projections of the ATR.

#### CST

In our protocol, we selected only the CST projections terminating in the precentral gyrus, postcentral gyri, and the posterior third of the superior frontal gyrus (SMA), as described previously (Chenot et al. 2019). We are aware that the CST includes additional axonal projections to more frontal regions (Dum et al., 2002). However, these were represented by fewer and sparser streamlines in our tractography data, and we thus decided to not include them in the present atlas. These more frontal CST projections may be harder for tractography to reconstruct consistently given their bending and fanning geometry, as opposed to the more straightforward CST projections to the motor regions. Future work exploring specific regions of interest for tractography seeding (e.g., the subthalamic nucleus) might help improve these results.

#### FX

The FX is a small bundle with high curvature throughout its extension. Its location in proximity of the ventricles makes it sensitive to partial voluming with CSF voxels (Vos et al 2011). These characteristics have made this bundle extremely challenging for tractography. To alleviate these limitations, we deployed a MSMT tractography algorithm (Jeurissen et al. 2014), which helped reduce the partial volume effect. We also avoided the use of constraining binary masks (WM, GM, CSF), which reduced the number of false negatives in the reconstructions. This approach allowed us to reconstruct the entire extent of the FX in most of the subjects. However, despite the successful reconstruction of this bundle in most subjects, a few reconstructions showed very few correct streamlines, and not all the subjects presented terminations extending into the temporal regions anterior to the hippocampus.

#### UF

The UF has been well-characterized in tractography studies. Although tractography is able to delineate the main trunk of the UF, it remains difficult to define its projections precisely and to separate them from those of the EmC, given their overlap. In our protocol, we aimed specifically at distinguishing these two projection systems, by including a ROI to separate the medial projections of the UF from the more lateral projections of the EmC (Heide et al 2013). We acknowledge the difficulty of completing this task accurately, as in most subjects it led to a reduced amount of UF streamlines reaching the superior frontal regions, with respect to those reaching the medial orbito-frontal regions.

#### SLF1

The exact human morphology of the SLF1 remains controversial, and its tractography-based reconstruction challenging, with inconsistent results (Wang et al. 2016). Particularly, while the literature overall agrees on its posterior terminations in the superior parietal lobule and precuneus, it remains unclear whether the anterior terminations of the SLF1 extend anteriorly to connect regions in the SFG and possibly cingulate cortex (Howells et al. 2018, Thiebaut de Schotten et al. 2012, Makris et al. 2005, Kamali et al. 2014), as observed in monkeys (Schmahmann et al. 2006, Thiebaut de Schotten et al. 2012), or whether they are constrained to the rostral part of the supplementary motor area (SMA) and pre-SMA (Hecht et al. 2015, Wassermann et al. 2013, Jang et al. 2012). This controversy arises from the fact that some previously published tractography studies could not reconstruct the most anterior streamlines of the SLF1 (Wassermann et al. 2013, Jang et al. 2012). While this might reflect a true inter-species difference, it might also be a tractography error due to the location of these fibers. They lie just underneath the u-shaped fibers of the SFG, in very close proximity to the CB, and at the intersection with major inferior-superior projection systems (CST and Corona Radiata) and the lateral projections of the CC. For the virtual dissection of the SLFI we adopted a protocol similar to what previously described by Howell et al. 2008 and we could recover the most frontal projections of the SLF1 in most of the subjects (Howells et al. 2018, Thiebaut de Schotten et al. 2012). However, even in these high-quality data, some subjects showed only few streamlines in this most frontal region, and a few subjects showed no streamlines at all. Future studies aimed at post-mortem validation of the anatomy of the frontal SLF1 will help elucidate whether this is due to the anatomical configuration and anatomical variability of this pathway or due to limitations of in vivo dMRI data (Maffei et al. 2020).

### 4.2 Automated reconstruction

We compared two ways in which our manual annotation protocol could be deployed for automated tractography: *(i)* Use the manually annotated streamlines to compute the anatomical neighborhood priors in TRACULA, and *(ii)* Use the manually defined ROIs as *post hoc* constraints in a multi-ROI method. We evaluated the accuracy of bundles reconstructed automatically with each approach, by comparing them to the manually annotated bundles in the same subject. We found that TRACULA achieved higher sensitivity (TPR) for the same reconstruction error (MHD), both overall (Fig. 8) and in individual bundles (Figs. 9-11). When comparing the multi-ROI method at its lowest reconstruction error and TRACULA at the same reconstruction error, the sensitivity achieved by TRACULA was an order of magnitude higher (Fig. 12). Performance gains with TRACULA were similar for association, commissural, and projection pathways (Fig. 13). Its performance was invariant to the inter-subject registration method (Fig. 14). Finally, when compared at the same level of sensitivity, the test-retest reliability of along-tract profiles extracted from microstructural measures was approximately four times greater for TRACULA than the multi-ROI approach (Fig. 15).

These performance differences may seem surprising, especially given that TRACULA is sometimes lumped together with multi-ROI methods in the literature. However, they can be explained by two fundamental algorithmic differences. First, multi-ROI methods typically use local tractography, which is prone to stopping or taking the wrong turn when it goes through challenging areas with complex fiber configurations. The role of the ROIs in a multi-ROI method is to remove these erroneous streamlines, but there is no guarantee that any correct streamlines will be left. The global tractography used by TRACULA models the complete trajectory of a bundle between its termination regions as a parametric curve. Thus, it is not possible for the paths generated by TRACULA to stop half-way between the regions.

The other key algorithmic difference is in how each method incorporates prior knowledge on the anatomy of the pathways of interest. Multi-ROI methods contain information about a set of regions that the pathway goes through, in template space coordinates. These regions are typically few (2-3), distant from each other, and deterministic. The anatomical neighborhood priors used by TRACULA contain detailed probabilistic information on how likely the pathway is to go through or next to each of the labels in a whole-brain anatomical segmentation. This information is encoded for anatomical neighbors in multiple directions and at multiple points along the pathway. These anatomical neighborhood priors implement the same idea as the “Markov priors” used in the FreeSurfer automated subcortical segmentation and cortical parcellation (Fischl et al. 2002; Fischl et al. 2004). The difference is that TRACULA uses the anatomical neighborhood priors to generate streamlines, not to classify voxels.

The fact that TRACULA relies on a structural segmentation from a T1-weighted scan may be viewed as a limitation. However, we have previously shown that TRACULA is robust to errors in the boundaries of the structural segmentation labels, or even to using a segmentation mapped from a different subject (Zöllei et al. 2019). That is because TRACULA only uses information on the relative position of WM pathways and structural segmentation labels (e.g., how frequently is pathway A medial to structure B), and not on their exact spatial coordinates. Furthermore, we have recently shown that it is possible to infer the full set of FreeSurfer segmentation and parcellation labels from a dMRI scan using deep learning (Ewert et al. 2020). Thus, a low-quality or missing T1-weighted scan is not an insurmountable problem.

A possible limitation of this study is that we did not compare TRACULA to *all* possible multi-ROI methods. However, we compared it to the manual annotation, which represents the best-case scenario of multi-ROI performance. The manually annotated bundles were generated from the b_max_=10,000 *s*/*mm*^2^ Connectom data, using state-of-the-art orientation reconstruction and probabilistic tractography techniques, augmented by painstaking manual editing. The bundles reconstructed automatically by TRACULA from b=1,000 *s*/*mm*^2^ data exhibited high sensitivity and low reconstruction error with respect to the manually annotated bundles. In addition, we compared TRACULA to a multi-ROI method that was automated and used the same input data and the same orientation reconstruction method as TRACULA. In that comparison, TRACULA exhibited much higher accuracy and reliability. In the future, it is possible to incorporate orientation reconstruction methods other than the ball-and-stick model in TRACULA.

Finally, our results demonstrate that tract-of-interest reconstruction, where the task is to reconstruct certain well-known, anatomically defined bundles, does not require a sophisticated dMRI acquisition protocol. Our automated reconstructions from b=1,000 *s*/*mm*^2^, 64-direction data achieved an overall sensitivity of 89% with respect to the manual annotations from b_max_=10,000 *s*/*mm*^2^, 512-direction data, for a reconstruction error of 3.5 *mm* for TRACULA and 4.2 *mm* for the multi-ROI method. Therefore, when the main use of dMRI data in a study is to reconstruct tracts of interest and analyze microstructural measures along them, the sophistication of the dMRI acquisition protocol should be determined by the microstructural measures and not by the tractography itself.

## 5. Conclusion

We have illustrated that TRACULA can take advantage of limited-availability, high-quality data that can only be acquired on a handful of Connectom scanners worldwide, to reconstruct white-matter bundles with high accuracy from more modest and widely available dMRI data. This allows the technological innovations of the HCP to benefit the wider community that does not have access to Connectom-style scanners. Both our WM tract atlas, which was annotated manually from Connectom data, and the software tools that can use it to reconstruct WM bundles in routine-quality data, are freely available as part of FreeSurfer 7.2.

